# Inhibition of oogenic JNK preserves fertility and ovarian hormones during DNA-damaging cancer therapy

**DOI:** 10.64898/2026.04.28.721450

**Authors:** Wenlong Zhao, Jiyang Zhang, Yingnan Bo, Yingzheng Wang, Mi Ran Choi, Shichao Liu, Qiang Zhang, So-Youn Kim, Shuo Xiao

## Abstract

Primary ovarian insufficiency (POI) and related infertility, early menopause, and endocrine disorders due to hormonal deficiency are major side effects in young female cancer patients undergoing cancer therapy. Current strategies preserving the fertility and hormonal functions of the ovary remain imperfect due to concerns of feasibility, efficacy, or safety. Herein, we identified c-Jun N-terminal kinase (JNK) as a pivotal regulator of the DNA damage response (DDR) signaling in oocytes of primordial follicles in response to DNA-damaging cancer therapy. Using pharmacological JNK inhibition and a genetically modified mouse model with oocyte-specific JNK deletion, together with histological, bioinformatic, and molecular approaches, we demonstrated that JNK inhibition prevented chemotherapy-induced oocyte apoptosis and POI, and preserved long-term reproductive cycles and fertility. Mechanistically, JNK was activated in response to chemotherapy-induced DNA damage in oocytes of primordial follicles, causing activation of transcription factor TAp63α and subsequent oocyte apoptosis, ultimately resulting in diminished ovarian reserve and POI. A more clinically relevant breast cancer-bearing mouse model revealed that JNK inhibition preserved the ovarian reserve without compromising anti-cancer efficacy of chemotherapy. Together, our study identifies oocyte-intrinsic JNK as a promising target for developing ovarian protectants and safeguarding reproductive health and fertility in young female cancer survivors.

## Introduction

Advances in cancer survival rates have greatly increased the awareness of the side effects of cancer therapy and long-term quality of life in cancer survivors, including primary ovarian insufficiency (POI) in childhood, adolescent, and young adult female cancer patients (1-3). Chemotherapy, along with irradiation, is gonadotoxic to the ovary, dramatically heightening the risks of POI and related delay or absence of puberty, infertility, early menopause, and endocrine disorders due to hormonal deficiency (4, 5). Women with POI also face other serious health consequences, including depression, osteoporosis, autoimmune disorders, heart disease, and increased mortality (6, 7).

The ovary contains follicles at various stages as its functional unit. Primordial follicles, the earliest stage of follicles in the ovary, consist of a central oocyte surrounded by a single flattened layer of granulosa cells. Primordial follicles are finite and non-renewable after birth, establishing a woman’s ovarian reserve that determines her reproductive lifespan. Depletion of the ovarian reserve leads to menopause. Oocytes within primordial follicles are particularly vulnerable to genotoxic insults and rely on a DNA damage response (DDR) signaling pathway to promptly detect and repair DNA lesions, or trigger apoptosis if the damage is irreparable (8-11). This highly evolved DDR ensures that only oocytes with a largely intact genome progress to ovulation and fertilization, preventing the transmission of genetic errors to the next generations. Most anti-cancer agents exert their effects by inducing DNA damage in highly proliferating cancer cells (12). However, these agents inadvertently cause DNA damage in the oocytes of primordial follicles, leading to their loss and associated POI, infertility, and hormonal deficiency as major off-target effects.

Current strategies preserving the ovarian reserve, fertility, and hormonal functions in young female cancer patients include oocyte/embryo cryopreservation, ovarian tissue cryopreservation (OTC), or ovarian suppression (13). These methods, however, remain imperfect due to concerns of feasibility, efficacy, or safety. Oocyte or embryo cryopreservation requires ovarian stimulation, which does not protect the ovarian reserve, delays cancer therapy, and is not feasible for prepubescent girls. OTC necessitates surgical removal of ovarian tissue, resulting in the loss of hormonal support of the ovary; subsequent ovarian tissue transplantation may reintroduce malignant cells (14). Ovarian suppression with gonadotropin-releasing hormone (GnRH) agonist remains controversial due to inconsistent efficacy and incomplete protection of the ovarian reserve (15, 16).

Developing ovarian protectants has emerged as a promising strategy for protecting and preserving the ovarian reserve in young female cancer patients. Such agents are applicable to both prepubescent girls and young adult women, do not require hormone stimulation or invasive surgery that delays cancer therapy, and safeguard both fertility and hormonal functions of the ovary (17, 18). Modulating key molecules within the DDR signaling pathway, such as the DNA damage-sensing kinases ATM or ATR, checkpoint kinases CHEK1 and CHEK2 (19), transcription factor TAp63α (20), and pro-apoptosis proteins NOXA and PUMA (10, 21), has been shown to prevent oocyte apoptosis induced by gonadotoxic anti-cancer agents. However, as the DDR signaling is largely shared between oocytes and cancer cells, these interventions risk compromising the efficacy of cancer therapy. Thus far, the precise mechanisms of DDR in oocytes, particularly how they differ from those in cancer cells, remain poorly understood. Therefore, it is critical to identify novel oocyte-intrinsic targets to preserve the ovarian reserve and fertility without interfering with the efficacy of cancer therapy in young female cancer patients.

In this study, we investigated the role of c-Jun NH2-terminal kinase (JNK), a stress-sensing kinase outside the canonical DDR cascade, in regulating the DDR signaling in oocytes of primordial follicles. We discovered that widely used DNA-damaging anti-cancer agents, including doxorubicin (DOXO), cisplatin (CDDP), and cyclophosphamide (CPA), prominently activated JNK in the oocytes and to a much lesser extent in granulosa cells of primordial follicles. Using a pharmacological JNK inhibitor and a genetically modified mouse model with oocyte-specific JNK deletion, together with histological, bioinformatic, molecular, and computational approaches, we demonstrated that JNK inhibition blocked chemotherapy-induced apoptosis in oocytes of primordial follicles and prevented POI, infertility, and other reproductive defects. Mechanistically, inhibition of JNK blocked the activation of TAp63α and downstream apoptotic signaling in oocytes of primordial follicles. Moreover, in a more clinically relevant breast cancer-bearing mouse model, JNK inhibition protected the ovarian reserve without compromising the anti-cancer efficacy of chemotherapy. Taken together, these results highlight oocyte-intrinsic JNK signaling as a promising target for the development of ovarian protectant, safeguarding reproductive health and fertility in young female cancer survivors.

## Results

### DNA-damaging chemotherapy induces POI and distinct transcriptomic changes between oocytes and granulosa cells of primordial follicles

To determine the gonadotoxic effect of chemotherapeutic agents, postnatal day 5 (PND5) female mice were treated with clinically relevant doses of 10 mg/kg DOXO, 7.5 mg/kg CDDP, and 150 mg/kg CPA once through intraperitoneal (i.p.) injection (Fig. 1A). On day 3-post treatment, histological staining and follicle counting revealed that DOXO, CDDP, or CPA depleted over 90% of the primordial follicles (92.25%, 91.82%, and 93.64%, respectively); primary follicles tended to decrease but the differences were not statistically significant; and secondary follicles were significantly reduced in all three chemo-groups (Fig. 1B). These results confirm the chemotherapy-induced POI. Our prior studies show that DNA-damaging chemotherapy activates DDR in oocytes of primordial follicles to induce oocyte apoptosis and primordial follicle death (22, 23). To identify key genes and signaling pathways involved in oogenic DDR, PND5 mice were treated with vehicle or 10 mg/kg DOXO via i.p.. Single oocytes and granulosa cells were collected from primordial follicles at 6 hours for single-cell SMART-seq2 RNA-seq analysis.

**Figure 1.**
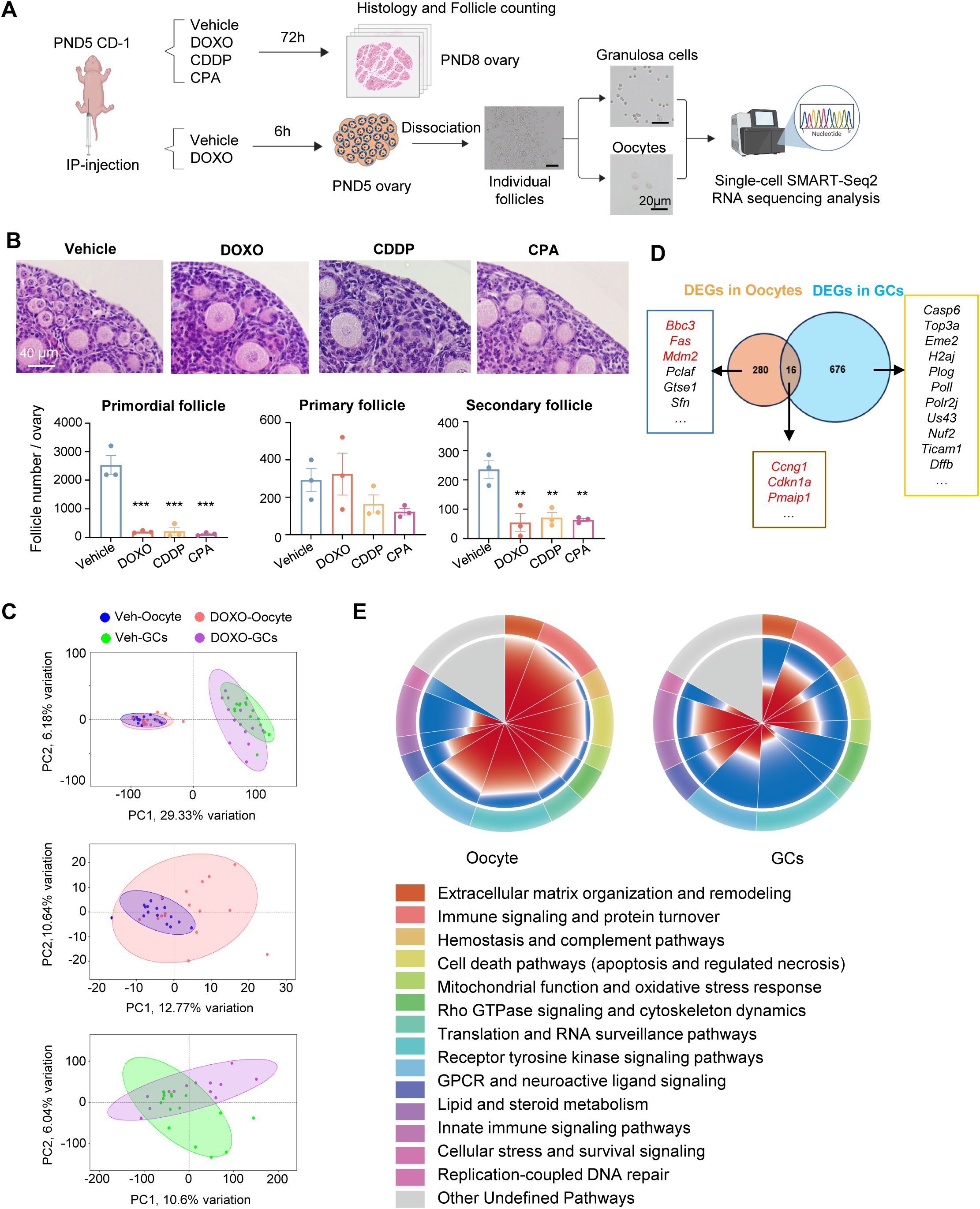
Gonadotoxic chemotherapy induces POI and distinct transcriptomic changes between oocytes and granulosa cells of primordial follicles. **(A)** Scheme of study design of chemotherapeutic treatment and endpoints. **(B)** Follicle counting with representative H&E staining of ovarian sections collected on day 3 post-treatment with vehicle, DOXO, CDDP, or CPA. (n=3 mice per group). Scale bar, 40 μm. Lower panels: quantification of primordial, primary, and secondary follicles per ovary in each treatment group. Data are presented as the mean ± SEM. One-way ANOVA followed by Tukey’s multiple comparisons test; ns, *p* > *0.05*; *, *p* < *0.05*; **, *p* < *0.01*; ***, *p* < *0.001*. **(C)** PCA of single-cell SMART-seq2 data from control and DOXO-treated oocytes (blue and red) and granulosa cells (GCs; green and purple). Each point represents an individual cell (n=16 oocytes per group, n=16 and 13 GCs in the Vehicle and DOXO groups, respectively). **(D)** Venn diagram of DEGs in oocytes and somatic cells after DOXO exposure. **(E)** GSEA on DEGs in oocytes (Left) and granulosa cells (Right), respectively. The outer ring represents the relative weight of each module (sum of absolute NES), while the inner ring shows the proportion of negatively (blue) and positively (red) enriched pathways compared to controls in the corresponding module.

Principal component analysis (PCA) separated oocytes and granulosa cells into two distinct clusters regardless of the treatment (Fig. 1C, upper panel), indicating distinct transcriptomes between the two cell types. Both oocytes and, particularly granulosa cells, exhibited a transcriptomic shift following DOXO treatment (Fig. 1C, middle and bottom panels, respectively). Compared to controls, there were 296 differentially expressed genes (DEGs) in oocytes and 692 DEGs in granulosa cells from DOXO-treated mice (Fig. S1B and Suppl. Table 2). Notably, only 16 DEGs overlapped between oocytes and granulosa cells, including DDR marker genes such as *Ccng1*, *Cdkn1a* (also termed *p21*), and *Pmaip1* (also termed *Noxa*) (Fig. 1D). Among 280 DEGs specifically altered in the oocytes, *Bbc3* (also termed *Puma*), *Fas*, and *Mdm2* are involved in DDR (Fig. 1D), and the deletion of *Bbc3* or *Puma* has been shown to protect the ovarian reserve during DNA-damaging chemotherapy or irradiation (10, 21).

Gene set enrichment analysis (GSEA) was performed based on differential gene expression between DOXO and control groups in each cell type using the Reactome gene set collection. Eight pathways were significantly enriched in the oocytes, whereas no pathways were enriched in the granulosa cells (Suppl. Table 3). The oocyte-enriched pathways were predominantly associated with translational regulation and RNA surveillance, indicating that oocytes had initiated an early stress response following DOXO treatment. To further compare global response patterns, we selected the top 200 pathways ranked by absolute normalized enrichment score (|NES|) in each cell type (Suppl. Table 3), constructed their union, and clustered them into functional modules. In the visualization shown in Fig. 1E, the outer ring represents the relative weight of each module (sum of |NES| values), while the inner ring shows the proportion of pathways with positive (red) or negative (blue) NES values compared to the control. Although the overall module weight distribution was similar between oocytes and granulosa cells, their positive/negative response patterns differed markedly in directionality.

In oocytes, multiple modules exhibited strongly coordinated responses following DOXO treatment. Using a stringent threshold to define the directional bias (>80% or <20% of pathways showing positive enrichment), 8 out of 12 modules showed strong activation, including receptor tyrosine kinase signaling pathways (80.85%), immune signaling and protein turnover (94.44%), cell death pathways (92.00%), extracellular matrix organization and remodeling (100.00%), Rho GTPase signaling and cytoskeleton dynamics (88.24%), translation and RNA surveillance pathways (85.71%), hemostasis and complement pathways (92.31%), and mitochondrial function and oxidative stress response (91.67%). Notably, the replication-coupled DNA repair module exhibited complete suppression, with 0.00% of pathways positively enriched and 100.00% negatively enriched. Collectively, these modules are primarily associated with stress signaling, immune activation, translational regulation, cytoskeletal remodeling, metabolic adaptation, and cell death, and their highly coordinated directional changes indicate that oocytes rapidly initiate a global, programmatic response that integrates stress sensing, proteostasis regulation, and cell fate determination following the treatment of DNA damaging chemotherapy. Different from oocytes, granulosa cells did not exhibit such globally coordinated responses. No modules showed strong activation bias, and only two modules, Rho GTPase signaling and cytoskeleton dynamics (11.76%) and translation and RNA surveillance pathways (14.29%), exhibited a strong suppression. Collectively, these results indicate that oocytes mount a highly coordinated, programmatic response to DOXO-induced stress than granulosa cells of primordial follicles.

### DNA-damaging chemotherapy activates JNK in oocytes of primordial follicles

Gene Ontology (GO) term enrichment analysis was next performed to identify key signaling pathways related to DEGs (Fig. 2A; Suppl. Table 4). DEGs overlapping between oocytes and granulosa cells are involved in cell cycle, apoptosis, reactive oxygen species (ROS), and autophagy. DEGs specifically altered in granulosa cells are related to tyrosine kinase phosphorylation, cell migration and division, endocytosis, and DNA damage. DEGs specifically changed in oocytes are primarily associated with protein transport, inflammation, proteolysis, hypoxia, apoptosis, and positive regulation of JNK cascade.

**Figure 2.**
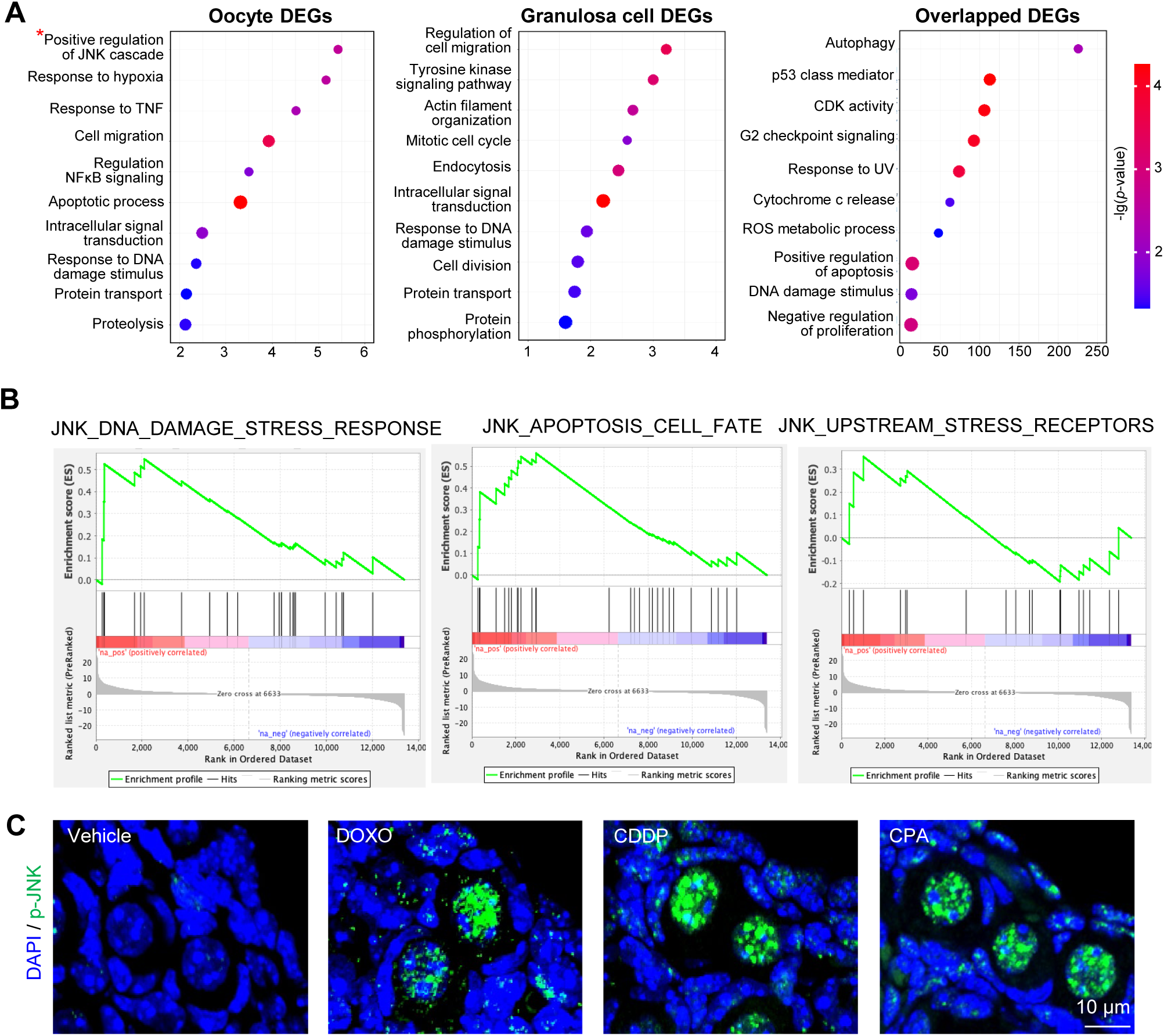
DOXO activates JNK in oocytes of primordial follicles. **(A)** GO biological process enrichment of overlapping, oocyte-specific, and granulosa-specific DEGs. **(B)** GSEA of oocyte gene profile on curated gene sets of JNK-related signaling pathways**. (C)** Immunofluorescence staining of ovarian sections indicating increased phosphorylated JNK (p-JNK, green) in primordial follicle oocytes 18 h after treatment with DOXO, CDDP, or CPA. Nuclei were counterstained with DAPI (blue). n=3 sections from 3 mice per group. Scale bar, 10 μm.

We next focused on JNK as it is selectively enriched in oocytes and acts as a central regulator of cell fate in response to diverse stress stimuli, including DNA damage (24-29). GSEA revealed a clear and coordinated activation of JNK-associated programs in oocytes following DOXO treatment (Fig. 2B, Fig. S2B, and Suppl. Table 5). In particular, gene sets representing stress-response and DNA damage–associated transcriptional programs were significantly enriched (NES=1.78, FDR=0.0042), indicating robust engagement of JNK-mediated stress signaling. In addition, gene sets associated with apoptotic and cell fate regulation also showed consistent positive enrichment (NES=1.61, FDR= 0.062), suggesting activation of downstream pathways that drive oocyte elimination. Other components of the JNK pathway, including upstream signaling modules and regulatory elements, exhibited concordant positive trends (NES=1.16-1.46, FDR=0.14-0.36), supporting a global activation of the JNK signaling. In contrast, granulosa cells did not exhibit significant enrichment of JNK-related gene sets (Fig. S2C and Suppl. Table 6).

Together, these bioinformatic data demonstrates that JNK signaling is selectively and robustly activated in oocytes of primordial follicles, particularly at the level of stress-response and apoptotic transcriptional programs, whereas granulosa cells fail to mount a coordinated JNK-mediated response. Indeed, JNK has also been implicated as pro-oncogenic in several tumor types, making it a widely pursued anti-cancer target (30-34). These features led us to hypothesize that JNK inhibition could be an advantageous ovarian protectant target that blocks oocyte death without compromising, or potentially even enhancing the efficacy of cancer therapy. Supporting this rationale, we next performed immunofluorescence and demonstrated that PND5 mice treated with DOXO, as well as other two widely used chemotherapeutic anti-cancer agents, CDDP and CPA, exhibited a robust increase in JNK phosphorylation, indicating JNK activation, in the oocytes of primordial follicles compared to surrounding somatic cells (Fig. 2C).

### Pharmacological inhibition of JNK prevents chemotherapy-induced POI

To determine the mechanistic role of JNK in chemotherapy-induced apoptosis in the oocytes of primordial follicles, PND5 female mice were treated with DOXO, CDDP, or CPA, with or without the co-administration of a selective JNK inhibitor SP600125 (Fig. 3A). DOXO, CDDP, or CPA alone depleted over 90% of the primordial follicles by day 3, confirming the induction of POI (Figs. 3B-D). SP600125 alone had no impact on follicle counts; however, it prevents the chemotherapy-induced loss of primordial follicles in all three chemo-groups (Figs. 3B-D). These results indicate that pharmacological inhibition of JNK protects oocytes from DNA-damaging chemotherapy-induced apoptosis, thereby protecting the ovarian reserve and preventing POI.

**Figure 3.**
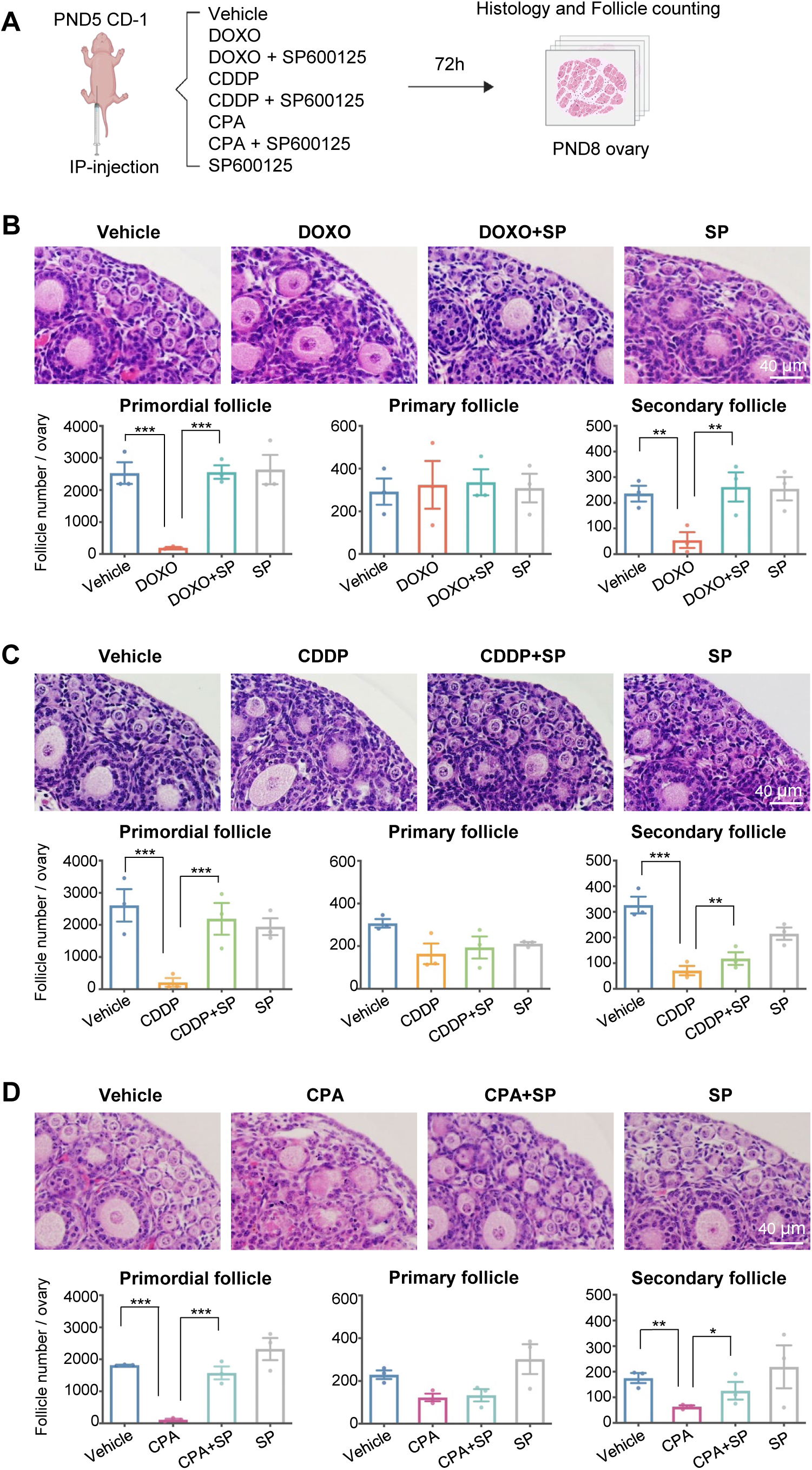
Pharmacological inhibition of JNK prevents chemotherapy-induced primary ovarian insufficiency (POI). **(A)** Experimental design of the POI mouse model induced by DOXO, CDDP or CPA, with or without the co-administration of JNK inhibitor SP600125. **(B-D)** Representative H&E staining of ovarian sections collected on day 3 post-treatment with DOXO (B), CDDP (C), or CPA (D), in the presence or absence of SP600125 (n=3 mice per group). Scale bar, 40 μm. Lower panels: quantification of primordial, primary, and secondary follicles per ovary in each treatment group. Data are presented as the mean ± SEM. One-way ANOVA followed by Tukey’s multiple comparisons test; ns, *p* > *0.05*; *, *p* < *0.05*; **, *p* < *0.01*; ***, *p* < *0.001*.

### Pharmacological inhibition of JNK preserves the ovarian reserve, ovarian cycles, and fertility long-term after DNA-damaging chemotherapy

We next applied the same chemotherapy regimen to assess if pharmacological JNK inhibition during gonadotoxic chemotherapy preserves the ovarian reserve, fertility, and other reproductive functions in long term (Fig. 4A). At 2 months post-treatment, DOXO, CDDP, or CPA alone significantly reduced the ovary weight (Fig. 4B) and depleted 84.36%, 87.37%, and 75.48% of primordial follicles, respectively (Figs. 4C and 4D). Similar depletion was observed at 8 months, with 80.95%, 74.21%, and 69.57% loss of primordial follicles, respectively (Figs. 4C and 4D). All three chemo-drugs also significantly reduced the counts of primary, secondary, and antral follicles at 2- and 8-month timepoints (Figs. 4C and 4D). SP600125 alone had no impact on the ovary weight, morphology, or follicle counts at all stages, but effectively prevented the reduction of the ovary weight and follicle counts across all developmental stages, even throughout the prolonged duration of 2 to 8 months post-chemotherapy (Figs. 4B-4D).

**Figure 4.**
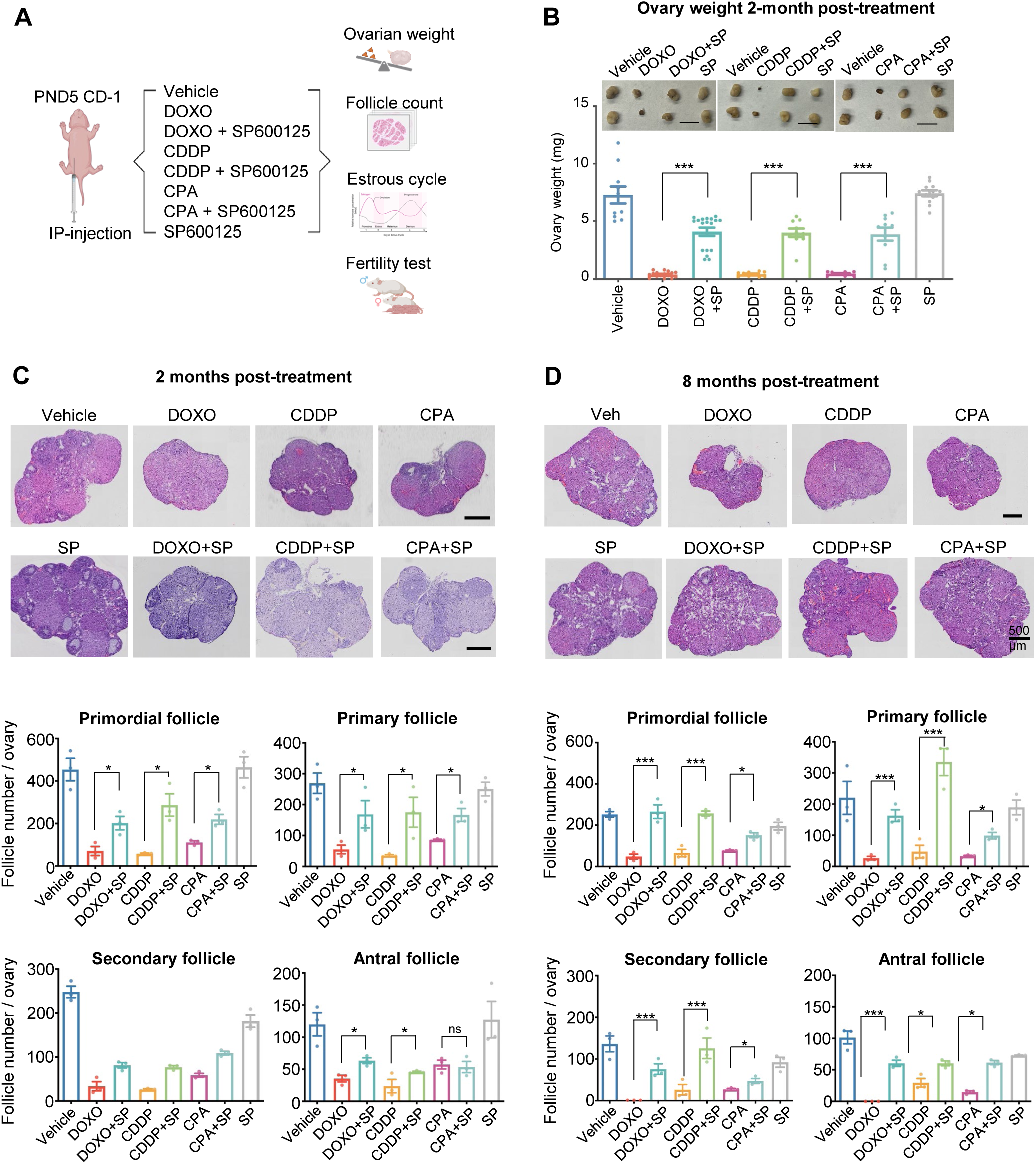
Pharmacological inhibition of JNK preserves the mouse ovarian reserve long-term after DNA-damaging chemotherapy. **(A)** Experimental design for examining the reproductive functions long term after DNA-damaging chemotherapy. **(B)** Representative images (upper) and statistical results (lower) of ovarian weight in mice treated with vehicle, chemotherapeutic agents, chemotherapeutic agents plus JNK inhibitor (SP600125), or JNK inhibitor alone at 2 months. Scale bar, 5 mm. **(C, D)** Representative ovarian histology from mice at 2 months (C, upper) and 8 months (D, upper) post-treatment with the indicated treatment regimens. n (mouse number with 2 ovaries per mouse) = 5 (Vehicle), 8 (DOXO), 9 (DOXO+SP), 5 (CDDP), 5 (CDDP+SP), 5 (CPA), 5 (CPA+SP), 5 (SP). Scale bar, 500 μm. Quantification of primordial, primary, secondary, and antral follicles at 2 months (C, lower) and 8 months (D, lower) post-treatment. n=3 ovaries from 3 mice per group. Data are presented as the mean ± SEM. Statistical significance was determined by one-way ANOVA; ns, *p* > *0.05*; *, *p* < *0.05*; **, *p* < *0.01*; ***, *p* < *0.001*.

The onset of mouse puberty, estrous cycles, and fertility were further assessed to evaluate whether JNK inhibition could rescue impaired reproductive functions caused by chemotherapy in long term. All three chemotherapeutic agents markedly delayed vaginal opening, with average opening day delayed to day 41, 39, and 40 in the DOXO-, CDDP-, and CPA-treated groups, respectively (Fig. 5A). Vaginal smears were performed for two consecutive weeks on mice at two months after treatment, which revealed disrupted estrous cyclicity in mice treated with all three chemo-drugs, evidenced by fewer days on proestrus/estrus and prolonged metestrus/diestrus (Figs. 5B and 5C). However, these reproductive defects were successfully prevented by JNK inhibitor (Figs. 5B and 5C).

**Figure 5.**
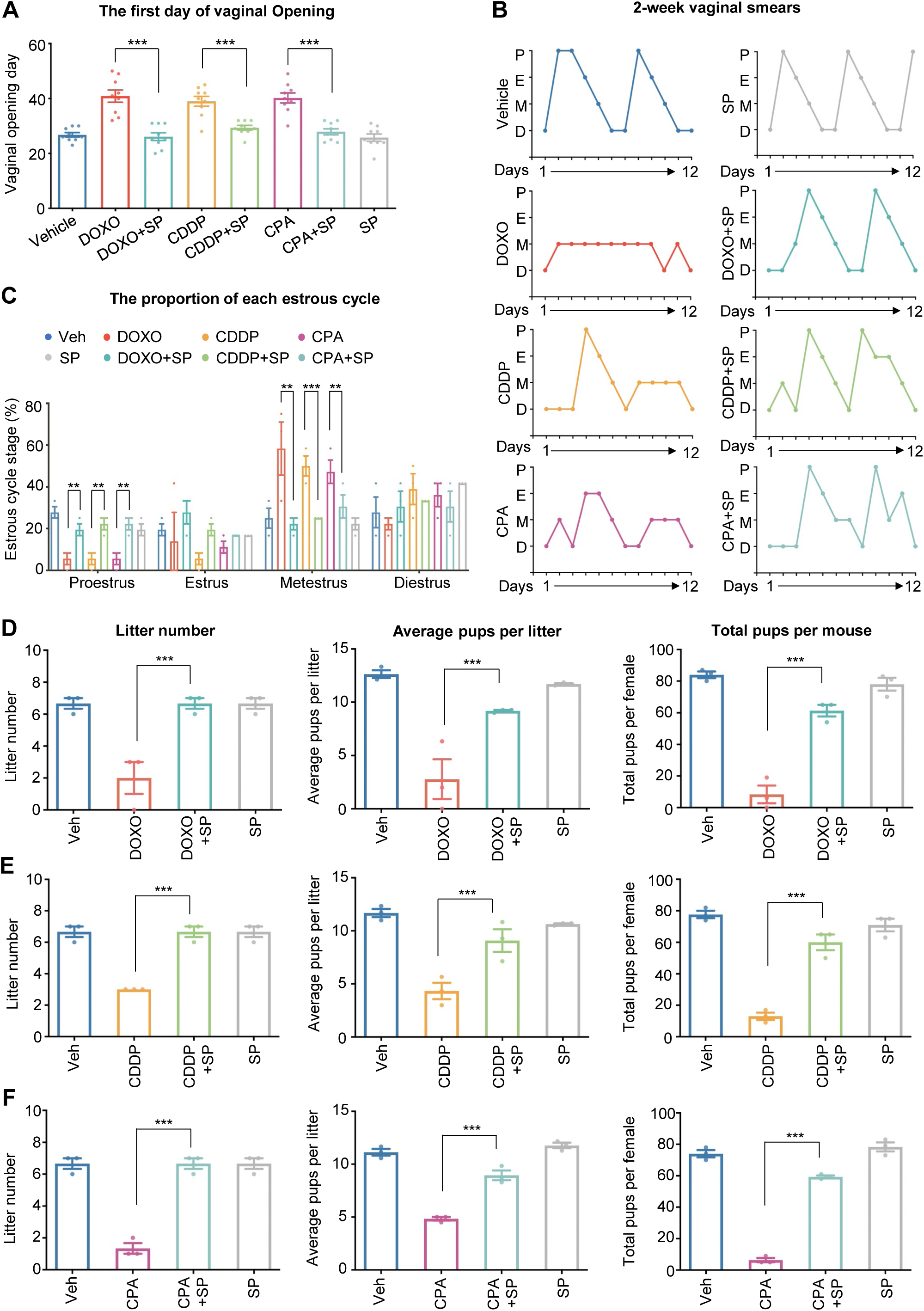
Pharmacological JNK inhibition preserves mouse puberty onset, estrous cycles, and fertility long-term after chemotherapy. **(A)** The timing of vaginal opening in mice treated with vehicle, chemotherapeutic agents (DOXO, CDDP, or CPA), chemotherapeutic agents plus the JNK inhibitor SP600125, or SP600125 alone (n=9 mice per group). **(B)** Representative 12-day vaginal smear profiles showing estrous cycles in 2-month-old mice treated with the indicated regimens. P, proestrus; E, estrus; M, metestrus; D, diestrus. **(C)** Quantification of the proportion of each estrous cycle stage across treatment groups (n=3 mice per group). **(D-F)** Litter number, litter size, and total pup numbers per mouse over a 5-month fertility test in various chemotherapeutic treatment groups (n=3 mice per group). Data are presented as the mean ± SEM. One-way ANOVA with Tukey’s correction. ns, *p* > *0.05*; *, *p* < *0.05*; **, *p* < *0.01*; ***, *p* < *0.001*.

A 5-month fertility test showed that mice treated with each of the chemotherapeutic drugs exhibited significantly reduced litter number, smaller litter size, and less total pups per animal, and mice became completely infertile at 3-5 months post-treatment (Figs. 5D-F). In contrast, mice receiving chemotherapy together with JNK inhibitor SP600125 maintained comparable fertility to the controls over a 5-month fertility test. Together, these results demonstrate that pharmacological inhibition of JNK preserves the mouse ovarian reserve, fertility, and related reproductive functions for long-term post-chemotherapy.

### Oocyte-specific deletion of JNK prevents chemotherapy-induced POI

Pharmacological experiments conducted above suggested that JNK contributes to chemotherapy-induced POI. However, whether this effect is mediated through oocytes, other ovarian cell types, or systemically needs to be confirmed. To address this, we generated a transgenic mouse model with oocyte-specific deletion of all three JNK genes, which avoids functional compensation between JNK isoforms. Single-cell RT-qPCR and immunofluorescence revealed significantly lower expression of *Jnk1-3* in oocytes, but not in ovarian somatic cells in *Jnk* cKO mice (Figs. S3A and 3B), confirming the oocyte-specific deletion of JNK. *Jnk* cKO mice displayed normal ovarian morphology and follicle counts (Figs. S4A and 4B), ovulation (Figs. S4C and 4D), and fertility (Figs. S4E and 4F), indicating that oogenic JNK deletion does not affect mouse ovarian functions and fertility.

To assess the oocyte-specific role of JNK in chemotherapy-induced oocyte apoptosis and POI, PND5 *Jnk^flox/flox^*and *Jnk* cKO mice were treated with DOXO, CDDP, or CPA as shown in Fig. 1A. By day 3, histology and follicle counting revealed 91.81%, 90.79%, 91.13% reduction of primordial follicles in DOXO, CDDP, and CPA-treated *Jnk^flox/flox^*mice, respectively, confirming chemotherapy-induced POI (Figs. 6A-6C). However, the lack of oogenic JNK in *Jnk* cKO mice successfully prevented primordial follicle depletion caused by all three chemotherapeutic agents (Figs. 6A-6C). Collectively, these results demonstrate the essential roles of oogenic JNK in regulating chemotherapy-induced apoptosis of primordial follicle oocyte and consequential POI.

**Figure 6.**
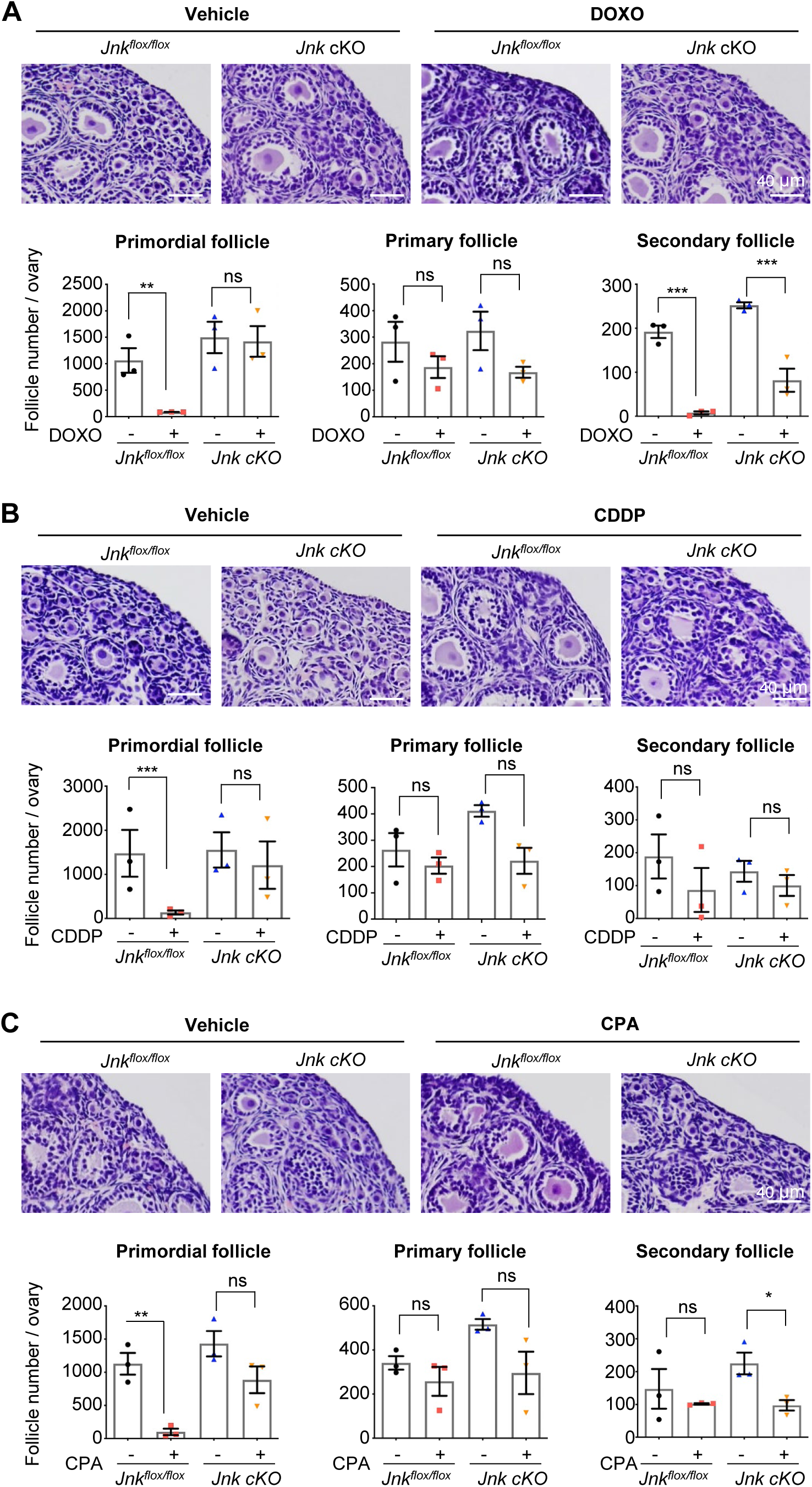
Oocyte-specific deletion of JNK prevents chemotherapy-induced POI. **(A, B, C)** Representative histological sections of ovaries from *Jnk^flox/flox^* and *Jnk cKO* mice collected 3 days after *in vivo* treatment with DOXO (A, upper), CDDP (B, upper) or CPA (C, upper). Scale bar, 40 μm. Quantification of primordial, primary, and secondary follicles (A-C, lower) in ovaries from *Jnk^flox/flox^* and *Jnk cKO* mice treated with vehicle or the indicated chemotherapeutic agents (n = 3 per group). Data are presented as the mean ± SEM. One-way ANOVA with Tukey correction. ns, *p* > *0.05*; *, *p* < *0.05*; **, *p* < *0.01*; ***, *p* < *0.001*.

### JNK regulates TAp63α activation in response to gonadotoxic chemotherapy

To further elucidate mechanistic roles of oogenic JNK, we collected mouse ovaries 6 hours post-treatment with DOXO, CDDP, or CPA, with or without co-administration of SP600125. Western blotting reveals a mobility shift of TAp63α, a p53 family protein transcription factor that critically determines the fate of DNA-damaged oocytes (20, 23, 35), in all three chemo-groups. This indicates hyper-phosphorylation and activation of TAp63α, which, however, was reversed by the co-administration of JNK inhibitor SP600125 (Figs. 7A). Ovaries from PND5 mice treated with vehicle or DOXO were collected for co-immunoprecipitation (Co-IP) analysis. TAp63α was detected in all input samples, with a mobility shift in DOXO-treated ovaries, indicating its hyper-phosphorylation and activation (Fig. 7B). TAp63α was precipitated with p-JNK antibody only in protein lysates from DOXO-treated ovaries, but not with IgG, nor in proteins from vehicle-treated ovaries (Fig. 7B), suggesting a physical interaction between activated JNK and TAp63α.

**Figure 7.**
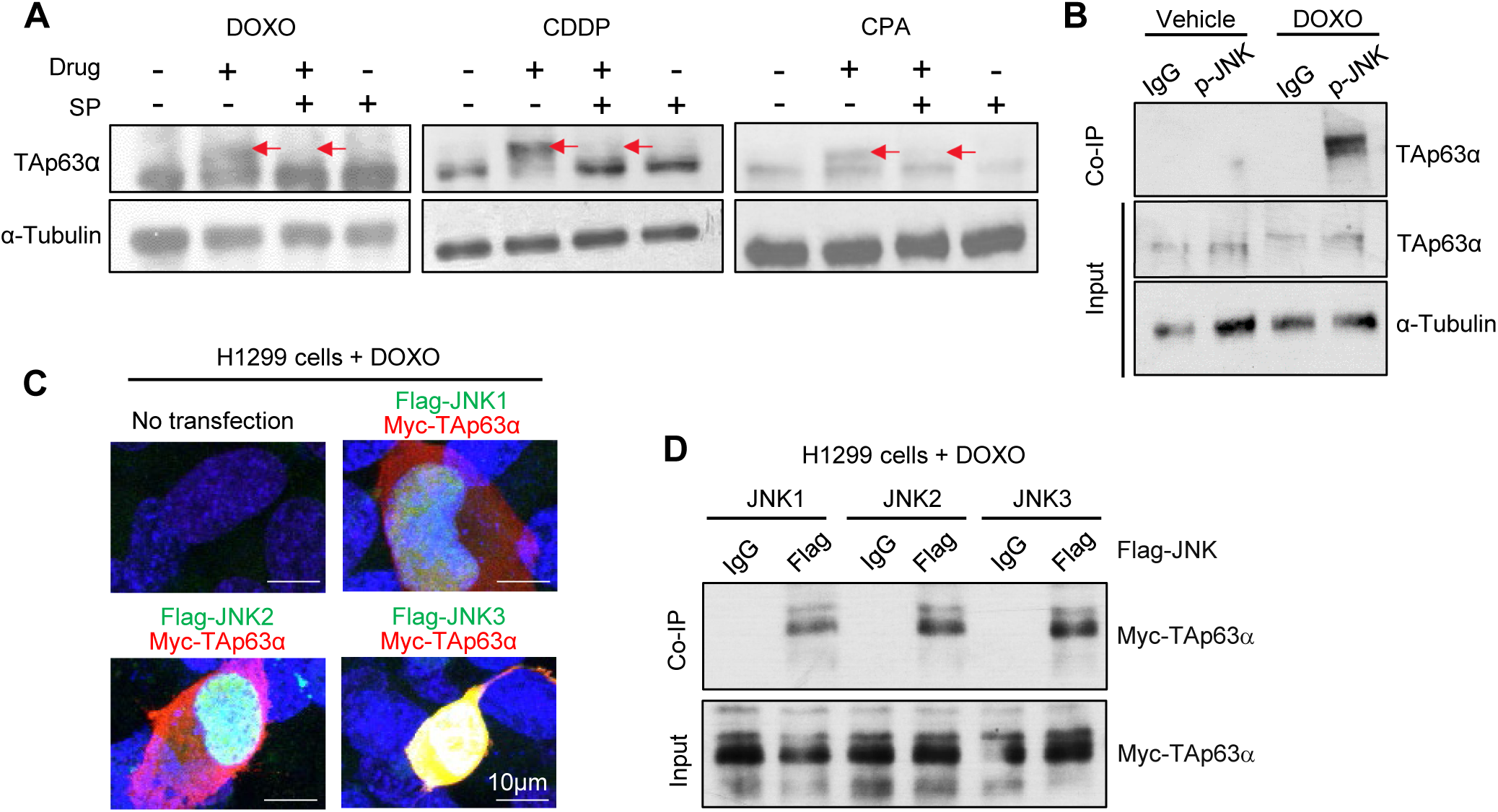
JNK regulates TAp63α activation following gonadotoxic chemotherapy. **(A)** Western blotting analyses showing the phosphorylation-dependent mobility shift of TAp63α in ovaries collected 18 h after treatment with DOXO (left), CDDP (middle), or CPA (right), with or without the co-treatment of JNK inhibitor SP600125. Red arrows indicate the hyperphosphorylated form of TAp63α. **(B)** Co-immunoprecipitation (Co-IP) with antibodies against p-JNK, indicating the interaction between p-JNK and TAp63α in ovaries from mice treated with DOXO for 18 h *in vivo*. **(C**) Immunofluorescence images, indicating the nuclear co-localization of Flag-JNK (green) and Myc-TAp63α (red) in H1299 cells treated with 2 μM DOXO for 24 hours. Scale bar, 10 μm. **(D)** Co-IP results indicating mechanistic interaction between JNK1/2/3 and TAp63α in H1299 cells treated with 2 μM DOXO treatment for 24 hours (n=3 replicates).

We next overexpressed JNK1/2/3 tagged with Flag and TAp63α tagged with Myc in H1299 cells to further investigate the protein-protein interaction (or binding) between JNK and TAp63α. Immunofluorescence staining revealed nuclear co-localization of Flag-JNK (green) with Myc-TAp63α (red) after treating the cells with 2 μM DOXO for 24 hours (Fig. 7C). Consistently, Co-IP using an anti-Flag antibody in cell lysates from H1299 cells treated with 2 μM DOXO for 24 hours showed the co-precipitated Myc–TAp63α with all three JNK isoforms in contrast to the Co-IP with IgG controls (Fig. 7D). Next, AlphaFold was used to predict the direct interaction between JNK and TAp63α. AlphaFold provides two key parameters: predicted Template Modeling (pTM) score and interface predicted Template Modeling (ipTM) score. A sum of pTM and ipTM > 0.5 indicates a high possibility of protein-protein interaction (29). The calculated sum of the AlphaFold scores were 0.88, 0.77, and 0.61 between TAp63 and JNK1, 2, and 3, respectively (Fig. S5), suggesting a high possibility of direct interaction between TAp63 and all three JNK isoforms. Together, the integration of *in vivo* and *in vitro* results with computational results highlight the direct protein-protein interaction between TAp63α and JNK.

### JNK inhibition preserves the ovarian reserve without compromising the anti-cancer efficacy of chemotherapy

To further explore the translational and therapeutic potential of JNK inhibition to preserve young cancer patients’ ovarian reserve and fertility, we established a syngeneic mouse breast cancer model using a mouse mammary carcinoma cell line, AT3OVA. (Fig. 8A). Tumors were allowed to grow to ∼ 50 mm³ and the mice were treated with DOXO or CPA, with or without JNK inhibitor SP600125. Tumor size in each experimental groups was measured at the timepoints of 0, 7, and 14 days post-treatment. Tumor volumes at day 7 and 14 were normalized to those at day 0. In vehicle controls, tumor volumes increased to 150-250 mm³ by day 7 and 200-350 mm³ by day 14 (Fig. 8B). Treatment with DOXO and CPA significantly reduced tumor volumes by 50.38% and 18.55% (DOXO) and by 55.20% and 24.42% (CPA) on day 7 and 14, respectively (Fig. 8B). Concurrently, both DOXO and CPA diminished the mouse ovarian reserve, evidenced by 80.3% and 74.25% loss of primordial follicles by day 14 in DOXO- and CPA-treated mice, respectively (Figs. 8C and 8D). JNK inhibitor alone did not affect tumor growth (Figs. 8B) nor the counts of all stages of follicles (Figs. 8C and 8D); however, it effectively prevented primordial follicle loss caused by DOXO and CPA (Figs. 8C and 8D). Taken together, using a translational breast cancer-bearing mouse model, recapitulating a condition of young female cancer patients, we demonstrate that JNK inhibition preserves the ovarian reserve without interfering with the anti-cancer efficacy.

**Figure 8.**
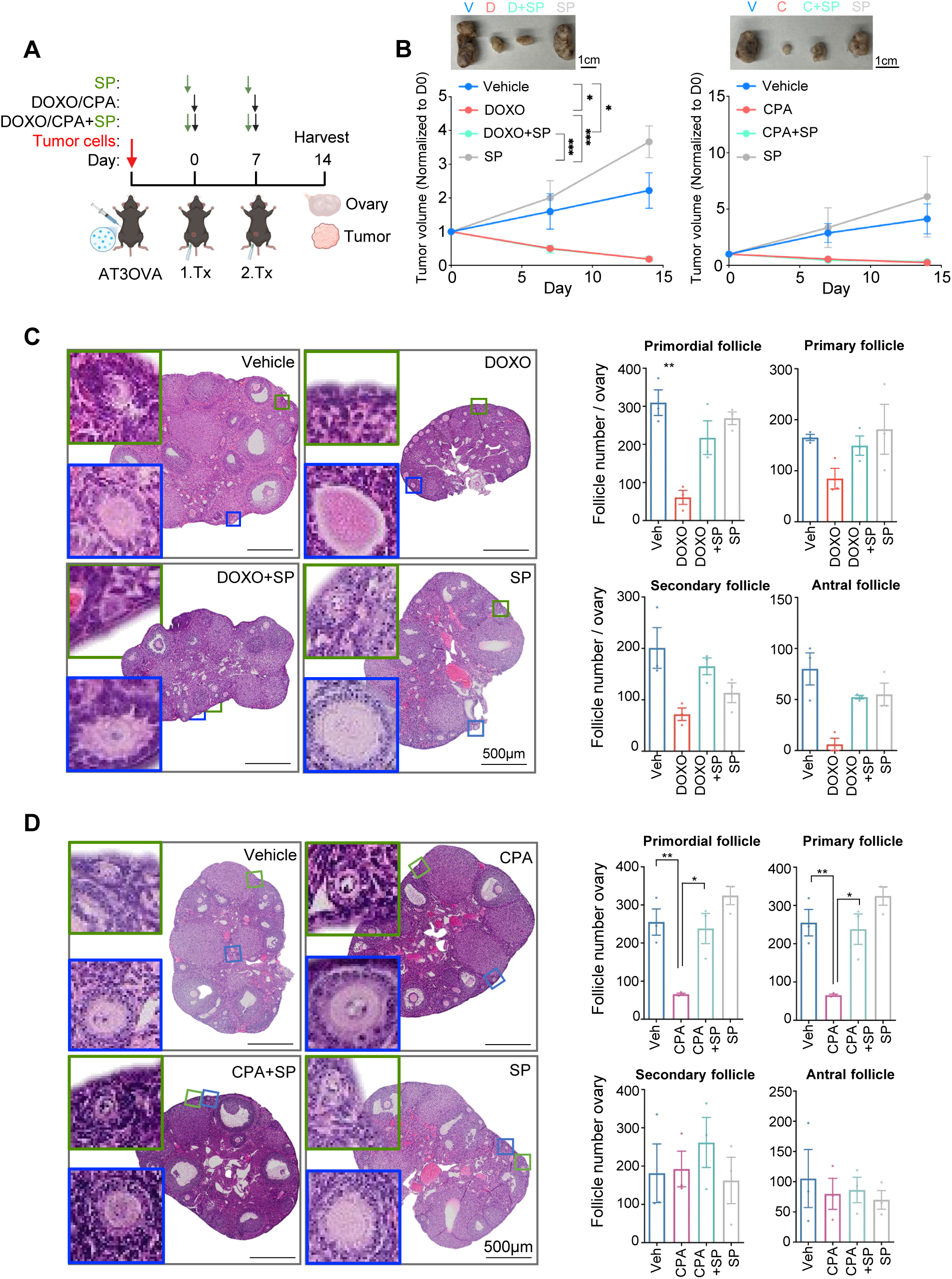
JNK inhibition preserves the ovarian reserve without compromising the efficacy of chemotherapy. **(A)** Experimental scheme. AT3OVA cells were orthotopically implanted into the mammary fat pad of adult female C57BL/6J mice. When tumors reached ∼50 mm³, mice received doxorubicin (DOXO, days 0 and 7) or a single dose of cyclophosphamide (CPA, 150 mg/kg, i.p.) with or without the JNK inhibitor SP600125 (10 mg/kg, i.p., 2 h prior to injection of chemo-agents). Tumors and ovaries were collected 7 days after the last treatment. **(B)** Tumor growth curves of AT3OVA tumor-bearing mice treated with vehicle, DOXO (left), CPA (right), SP600125, or in combination. Tumor size was normalized to the measurement taken on the day before the beginning of chemotherapy. The number of mice used for DOXO treatment; N=3 (Vehicle), 3 (SP600125), 4 (DOXO), and 4 (DOXO+SP600125). The number of the mice for the CPA treatment; N=4 (Vehicle), 4 (SP600125), 3 (CPA), and 3 (CPA+SP600125). Insets show representative tumors collected at day 14. Scale bar, 1 cm. **(C and D)** Representative histological images of ovarian sections from the mice treated with DOXO (C) and with CPA (D). Insets show primordial follicle (green box) and primary follicle (blue box). Scale bar, 500 μm. Quantification results of primordial, primary, secondary, and antral follicles per ovary in each group (n=3 mice per group) are shown for DOXO-treated group (C, right) and for CPA-treated group (D, right). Data represent mean ± SEM; One-way ANOVA with Turkey correction. ns, *p > 0.05*; *, *p < 0.05*; **, *p < 0.01*; ***, *p < 0.001*.

## Discussion

DNA-damaging cancer therapy activates DDR signaling pathway in the oocytes of primordial follicles, leading to apoptosis, depletion of ovarian reserve, and POI. Previous studies, including our own, demonstrated that inhibition of several key DDR molecules prevents oocyte apoptosis induced by gonadotoxic cancer therapy (10, 19-21). However, most of the DDR components are largely shared between oocytes and cancer cells, and, therefore, targeting these molecules raises concerns of interfering with the anti-cancer efficacy and/or compromising genome stability in somatic tissues. Here, we established JNK, as a non-canonical genotoxic sensing kinase, integrating cancer therapy-induced DNA damage to the activation of TAp63α and apoptosis in oocytes of primordial follicles, and JNK inhibition preserves the ovarian reserve without impairing anti-cancer efficacy.

Although all three chemotherapeutic agents used here induce DNA damage, their mechanisms differ. DOXO acts as a topoisomerase II (TOP2) inhibitor, blocking the resealing of DSBs during DNA replication (36). CDDP and the active metabolite of CPA generate intra- and inter-strand DNA crosslinks (37, 38). In oocytes of primordial follicles, DOXO activates ATM, which phosphorylates CHK2; together with casein kinase 1 (CK1), CHK2 promotes the hyper-phosphorylation of TAp63α, converting it from an inactive dimer to an active tetramer, triggering oocyte apoptosis (23, 39, 40). CDDP activates both ATR/CHK1 and ATM/CHK2 pathways to promote TAp63α-mediated oocyte apoptosis (10, 20, 40, 41). CPA has been shown to cause growing follicle atresia to exhaust the ovarian reserve (42-45), but accumulating evidence indicates that CPA also induces oocyte DNA damage through the ATR➔CHK1/CHK2➔TAp63α axis (21, 46-49). Our results show that both the pharmacological inhibition and a mouse model with oocyte-specific deletion of JNK prevents chemotherapy-induced hyperphosphorylation of TAp63α. Because Gdf9-iCre is active exclusively in the oocytes of primordial follicles, the protective phenotype observed in *Jnk* cKO mice reflects a direct and oocyte-intrinsic requirement for JNK signaling. Co-IP assay and AlphaFold analysis further demonstrate JNK is required for the activation of TAp63α and downstream apoptosis execution signaling. Although future studies are required to define whether JNK directly phosphorylates TAp63α or requires scaffolding or kinase crosstalk, these results highlight that unlike the canonical DDR kinases that directly sense DNA lesions, JNK is a crucial regulator sensing the genotoxic stress and controlling the activation of DDR signaling pathway and downstream apoptosis execution in the oocytes of primordial follicles.

Mammals express three JNK isoforms (50). JNK1 and JNK2 are broadly expressed, whereas JNK3 is enriched in neural and reproductive tissues (25, 50). JNK1 is the primary driver of stress-induced apoptosis, while JNK2 and JNK3 exhibit context-dependent or supporting roles (51, 52). SP600125 used here targets all three JNKs and our *Jnk* cKO mice have all *Jnk* alleles deleted, minimizing the compensatory effects. The Co-IP and AlphaFold results indicate interactions between TAp63α and all three JNK isoforms, suggesting collective contributions of JNK1, JNK2, and JNK3 to the activation of oogenic DDR signaling. Future studies using single- and dual-isoform *Jnk* knockout mouse models are warranted to dissect isoform-specific contributions to oocyte apoptosis.

JNK has been implicated to promote tumor growth, making JNK inhibition an attractive anti-cancer strategy (33, 52). Its pro-tumorigenic role in breast cancer is subtype specific. In HER2-positive tumors, JNK cross-talks with HER2/PI3K signaling pathway to promote survival and trastuzumab resistance (53). In triple-negative breast cancer (TNBC), JNK drives epithelial–mesenchymal transition (EMT), chemoresistance via autophagy, and immune evasion through PD-L1 upregulation and stromal remodeling (54, 55). In patient-derived xenograft (PDX) and E0771 syngeneic TNBC models, JNK inhibitor JNK-IN-8 enhances chemotherapy efficacy by reversing EMT phenotypes and reducing immune evasion through PD-L1 suppression (55). SP600125 has also been shown to inhibit TNBC invasiveness by blocking the JNK/c-Jun/MMP-9 axis (56) and to mitigate JNK/AP-1–mediated cell death following microtubule acetylation defects (57). Interestingly, in our TNBC-derived breast cancer mouse model, SP600125 neither reduced tumor size alone nor enhanced the anti-cancer efficacy of DOXO or CPA. This discrepancy from prior reports may arise from multiple factors, including intrinsic genetic differences between immortalized lines and PDX/E0771 models, tumor heterogeneity, pharmacodynamic differences between SP600125 and JNK-IN-8, and potential drug delivery limitations. Future studies are warranted to test ovarian-protective effects of JNK inhibition using clinically relevant breast cancer models as well as other malignancies common in young women, including hormone receptor–positive breast cancer, leukemia, and Hodgkin lymphoma.

The fate of DNA-damaged oocytes in primordial follicles is determined by a delicate balance between survival after DNA repair and elimination via apoptosis. Mice deficient in PUMA or NOXA retain nearly entire ovarian reserve and normal fertility, with offspring showing no congenital abnormalities or developmental deficits (10, 21). Consistent with this, JNK inhibition protects oocytes from apoptosis following genotoxic cancer therapy, thereby preserving the ovarian reserve, reproductive cycles, and fertility. However, whether damaged DNA persists during oogenesis remain unresolved. Future studies are needed to determine whether oocytes harbor unrepaired genomic lesions or if the DDR machinery fully restores genomic integrity to safeguard offspring health.

In conclusion, through bioinformatic, pharmacological, genetic, molecular, and computational approaches, we demonstrate for the first time that JNK is a crucial regulator contributing to the activation of DDR-TAp63α signaling pathway and apoptosis in the oocytes of primordial follicles. Inhibition of oogenic JNK prevents chemotherapy-induced oocyte apoptosis, POI, and infertility without compromising anti-cancer efficacy. These findings provide a promising therapeutic avenue to develop ovarian protectants that preserve the ovarian reserve, fertility, and endocrine function in young female cancer survivors undergoing gonadotoxic cancer therapy. Future effort will prioritize translational strategies, including patient-derived tumor xenografts and early-phase clinical trials, to validate JNK inhibition as an ovary-protective strategy. Further, defining isoform-specific JNK functions in cancer biology and oocyte preservation should guide the development of next-generation, highly selective JNK inhibitors with improved safety profiles.

## Materials and Methods

### Animals

Wild-type (WT) CD-1 and C57BL/6 mice were purchased from Envigo (Indianapolis, IN, USA). Mice with floxed alleles of *Jnk1*, *Jnk2*, and *Jnk3* were gift from Dr. Roger J. Davis (University of Massachusetts Medical School, Worcester, MA, USA). *Gdf9-icre* mice were gift from Dr. Karen Schindler (Rutgers University, NJ, USA). All animals were housed under constant temperature, humidity, and a 12-h light/dark cycle, with *ad libitum* access to food and water. All procedures on mice were approved by the Institutional Animal Care and Use Committee (IACUC) at Rutgers University.

To obtain female mice with oocyte-specific deletion of all three *Jnk* genes (*Jnk1/2/3^flox/flox^;Gdf9-icre^+^*, hereafter referred to as *Jnk1/2/3* conditional knockout or *Jnk* cKO), *Jnk1/2/3^flox/flox^*females were mated with *Jnk1/2/3^flox/flox^;Gdf9-icre^+^*males. Mice genotyped as *Jnk1/2/3^flox/flox^;Gdf9-icre^-^*(hereafter referred to as *Jnk^flox/flox^*) were used as controls. The sequences of gene primers for genotyping polymerase chain reactions are provided in Supplementary Table 1.

### Treatment with chemotherapeutic drugs

CD-1 female mice at postnatal day 5 (PND5) were treated with vehicle, 10 mg/kg doxorubicin (DOXO, 44583, Sigma-Aldrich), 7.5 mg/kg cisplatin (CDDP, P4394, Sigma-Aldrich), or 150 mg/kg cyclophosphamide (CPA, PHR1404, Sigma-Aldrich, St. Louis, MO, USA), respectively, via an one-time intraperitoneal (IP) injection to induce apoptosis in oocytes of primordial follicles, follicle atresia, and POI, as described previously (23, 46). To assess the role of JNK in chemotherapy-induced POI, PND5 females were injected with 10 mg/kg SP600125 (HY12041, MedChemExpress, Monmouth Junction, NJ, USA), a selective JNK inhibitor targeting all three JNK isoforms, via i.p. 2 hours before the administration of each chemotherapeutic agent. Ovaries were collected after 3 days, 2 months, or 8 months post-treatment for histology and follicle counting across all developmental stages (22, 23, 58).

### SMART-seq2-based single-cell RNA sequencing (scRNA-seq) and data analyses

Whole ovaries were harvested from PND5 CD-1 female mice (six ovaries in total) at 6 hours post-treatment with vehicle or 10 mg/kg DOXO via i.p. for single-follicular-cell SMART-seq2 RNA-seq analysis. Ovaries were cut into eight pieces and digested with 30.8 μg/mL Liberase TM (5401119001, Roche, Basel, Switzerland) and 456 IU/mL DNase I (LS002139, Worthington Biochemical Corporation, Lakewood, NJ, USA) in Leibovitz’s L-15 medium (11415064, Thermo Fisher Scientific, Waltham, MA, USA) at 37℃, 5% CO_2_ for 20 min. Partially digested tissues were further dissociated mechanically by repeated pipetting with a P1000 pipette.

The dissociated ovarian cells were collected by a brief centrifugation at 200 g, followed by an additional digestion with Accutase (A1110501, Thermo Fisher Scientific) for 5 min at 37℃ and in 5% CO_2_. The cell suspension was gently pipetted up and down for 10 times to obtain a single-cell suspension, which was then passed through a 100 μm- cell strainer to remove debris and tissue clumps. After centrifugation at 1000 rpm for 3 min, the supernatant was discarded, and the pellet was resuspended in PBS. Single somatic cells and oocytes were manually collected under a dissection microscope using a mouth-controlled glass pipette. Oocytes with a diameter of approximately 10 μm were classified as oocytes from primordial follicles. Because isolation of single follicular cells manually may inevitably capture a small number of non-follicular cells, cell identities were further verified at the transcriptomic level in the subsequent single-cell RNA-seq analysis.

The single-cell SMART-seq2 RNA-seq libraries were constructed using a modified single-cell tagged reverse transcription sequencing protocol as previously described (59). Briefly, mRNA from each single cell was extracted into the lysis buffer provided by the SMART-seq v4 Ultra-low input RNA kit (634891, TaKaRa, San Jose, CA, USA). mRNAs were then reverse-transcribed into complementary DNA (cDNA), which was then pre-amplified and purified. Single-cell transcriptomes were subsequently tagged with unique barcodes using the Nextera XT DNA library preparation kit (FC-131-1096, Illumina, San Diego, CA, USA) and the Nextera XT Index kit v2 setA (TG-131-2001, Illumina). Libraries were then pooled and sequenced on a NovaSeq PE150 (Illumina).

The raw sequencing of SMART-seq2 data were processed using the Partek Flow platform with default parameters to generate gene raw counts and identify differentially expressed genes (DEGs) between control and DOXO-treated groups. Only single cells with high oocyte and granulosa marker genes were used in the following bioinformatic analysis and comparison. Fig. S1A shows the selected marker genes of oocytes (*Ddx4, Lhx8,* and *Gdpd1*) and granulosa cells (*Foxl2, Arhgef17,* and *Socs3*) from primordial follicles that have been previously reported with single-cell RNA-seq studies (60, 61).

Standard principal component analysis (PCA) was performed separately for oocytes, granulosa cells, and for the combined dataset to assess overall transcriptomic differences. Gene Ontology (GO) term and Kyoto Encyclopedia of Genes and Genomes (KEGG) pathway enrichment analyses were conducted using online platform Heml 2.0 (https://hemi.biocuckoo.cn). Venn diagram was generated using the online tool Venny 2.1 (https://bioinfogp.cnb.csic.es/tools/venny/). R software (version 4.2.1) was used to generate the volcano plots by EnhancedVolcano package. Overlapped DEGs as well as top 20 up- and down-regulated genes in oocytes and granulosa cells were visualized by heatmap using heatmap package in R.

Gene set enrichment analysis (GSEA) was performed using the GSEA software (Broad Institute) in preranked mode. Differential gene expression results for oocytes and granulosa cells were used to generate ranked gene lists based on log2 fold change (DOXO-treated vs. control). Gene identifiers were converted to mouse gene symbols, and duplicate genes were removed by retaining the entry with the highest absolute log2 fold change. Custom JNK-related gene sets were constructed to represent distinct functional layers of the JNK signaling pathway, including upstream stress receptors, core MAPK cascade components, regulatory scaffolds, downstream AP-1 transcriptional targets, apoptosis-related effectors, and DNA damage–associated stress response genes. These gene sets were formatted in GMT format and used as the gene set database. GSEA was performed with 1000 permutations, using the gene set permutation type. No collapsing or remapping of gene identifiers was applied. Enrichment results were evaluated based on normalized enrichment score (NES), nominal p-value, and false discovery rate (FDR). Gene sets with FDR < 0.25 were considered significantly enriched. Leading-edge subsets were used to identify genes contributing most to the enrichment signal.

### Histology and follicle counting

Ovaries were fixed overnight in Modified Davidson’s fixative (64133-50, Electron Microscopy Sciences, PA, USA), dehydrated through a gradient ethanol series, embedded in paraffin, and serially sectioned at 5 μm thickness using an RM 2165 microtome (149030010, Leica Microsystems, Nussloch, Germany). All sections were stained with hematoxylin and eosin (H&E, STLSL9016 and 50420843, Thermo Fisher Scientific) following a protocol established in our previous studies (22, 23, 58, 62). Follicles were classified as primordial, primary, secondary, and antral follicles based on established morphological criteria. Every fifth ovary section was used for histological staining and follicle counting.

### Vaginal smears, estrous cycle determination, and fertility test

Estrous cyclicity was assessed in 2-month-old female mice by daily vaginal smears performed at approximately 14:00 for 12 consecutive days (covering 2-3 full estrous cycles). Briefly, the vaginal canal was gently flushed with PBS using a P200 pipette, and the collected fluid was smeared onto a glass slide for microscopic examination. The dominant cell type in vaginal cytology was used to determine the stage of the estrous cycle: Proestrus (P) - predominantly round, nucleated epithelial cells; Estrus (E) - predominantly cornified and anucleated squamous epithelial cells; metestrus (M) - a mixture of epithelial cells and leukocytes; Diestrus (D) - nucleated epithelial cells with a predominance of leukocytes. For fertility test, PND56 (8 weeks) female mice from each group were paired with fertility-proven CD-1 WT male mice. Fertility and litter size were monitored for 5 months.

### Quantitative real-time PCR (RT-qPCR)

To confirm the oogenic deletion of *Jnk*, single somatic cells and oocytes from primordial follicles were dissociated from PND5 *Jnk cKO* mice. Ovarian cell dissociation was performed using the same protocol described above for SMART-seq2. Total single-cell RNA was extracted using PicoPure™ RNA Isolation Kit (Kit0204, Thermo Fisher Scientific), followed by the synthesis of complementary DNA (cDNA) using the Superscript III first-strand Synthesis kit (18080051, Invitrogen, Carlsbad, CA, USA). Primer sequences for all genes are listed in Supplementary Table 1. RT-qPCR was performed using SYBR Green Master Mix (4367659, Applied Biosystems, Foster City, CA, USA). The housekeeping gene *Gapdh* was used as the internal control, and the comparative cycle threshold (ΔΔCT) method was applied for gene expression analysis. Fold-changes were calculated relative to the vehicle-treated samples.

### Western blotting

Ovaries were collected at 18 h post-treatment from the following groups: vehicle control, chemotherapeutic drug alone, co-administration of a chemotherapeutic drug (DOXO, CDDP, or CPA) and JNK inhibitor, or JNK inhibitor alone. Ovaries were homogenized in the tissue extraction buffer (FNN0071, Thermo Fisher Scientific) supplemented with protease inhibitor cocktail (87786, Thermo Fisher Scientific) and phosphatase inhibitor cocktails (78420, Thermo Fisher Scientific). Debris was removed through centrifugation at 12,000 g for 10 min at 4 ℃. Protein extraction was boiled at 95 ℃ for 5 min to denature proteins before loading into SDS-PAGE gel for electrophoresis. Subsequent procedures were carried out following the standard Western blot protocols. The primary antibodies used include: TAp63α (1:1000; 39692s, Cell Signaling Technology, Danvers, MA, USA), α/β-tubulin (1:5000; 2148s, Cell Signaling Technology), Myc (1:5000; 16286-1-AP, Proteintech, Rosemont, IL, USA) and Flag (1:5000; 66008-4-LG, Proteintech). Proteins were detected using Pierce^TM^ ECL plus Western blotting substrate kit (32132, Thermo Fisher Scientific) according to manufacturer’s instructions.

### Cell culture, transfection, and vector constructs

H1299 cells (CRL-5803, ATCC, Manassas, VA, USA) were cultured in RPMI-1640 (11875093, Thermo Fisher Scientific) supplemented with 10% fetal bovine serum (FBS). For transfection assays, H1299 cells were transfected with *Myc-TAp63*α and Flag-tagged *Jnk1*, *Jnk2*, or *Jnk3* using Lipofectamine^TM^ 2000 (18324012, Thermo Fisher Scientific) according to the manufacturer’s instructions. All expression vectors were custom constructed by VectorBuilder company (Chicago, IL, USA).

### Immunofluorescence microscopy

Immunofluorescence staining was performed as described in our previous studies (22, 58, 62). Briefly, ovary sections were deparaffinized in 100% xylene for 10 min and rehydrated in 100% ethanol for 6 min, followed by antigen retrieval using 10 mM sodium citrate buffer (pH 6.0). Sections were permeabilized in 0.2% Triton X-100 in PBS and blocked with 5% bovine serum albumin (BSA) in PBS for 1 h at room temperature. Primary antibodies were applied and incubated overnight at 4℃. Sections were then incubated with secondary antibodies for 1 h at room temperature, and nuclei were counterstained using Hoechst 33258 (94403, Thermo Fisher Scientific). Sections were mounted with proLong^TM^ Gold antifade mountant (P10144, Thermo Fisher Scientific).

H1299 cells transfected with *Myc-TAp63*α and Flag-tagged *Jnk1/2/3* were seeded on 12 mm diameter glass coverslips and treated with 2 μM DOXO for 24 h before fixation. Fixed H1299 cells were permeabilized and blocked in 0.2% Triton X-100 in PBS with 5% BSA for 1 h at room temperature. Incubations with primary and secondary antibodies were performed as described above. Antibodies used include: phospho-JNK (p-JNK; 1:1000; ab124956, Abcam Cambridge, UK), total JNK (1:1000; ab179461, Abcam), Myc (1:1000; 16286-1-AP, Proteintech, Rosemont, IL, USA), Flag (1:1000, 66008-4-LG, Proteintech), Alexa Fluor 488–conjugated goat anti-rabbit IgG (1:1000; A11008, Thermo Fisher Scientific), and Alexa Fluor 555–conjugated donkey anti-mouse IgG (1:1000; A10036, Thermo Fisher Scientific). Fluorescence signals were measured using Image J under the same exposure settings, with all experiments performed in triplicate.

### Co-immunoprecipitation (Co-IP)

H1299 cells transfected with *Myc-TAp63*α and Flag-tagged *Jnk1/2/3* were treated with 2 μM DOXO for 24 h and then harvested in Co-IP lysis buffer supplemented with protease inhibitor cocktail and phosphatase inhibitor cocktail. Ovaries from PND5 CD-1 mice with or without co-administration of DOXO were collected at 18 h for Co-IP. The lysate was centrifuged at 12,000 g for 10 min at 4℃ to remove debris, and supernatants were processed according to the manufacturer’s instructions for the Pierce™ Co-Immunoprecipitation Kit (26149, Thermo Fisher Scientific). Briefly, primary antibodies against Myc (1:1000; 16286-1-AP, Proteintech), flag (1:1000; 66008-4-LG, Proteintech), or p-JNK (1:1000; ab124956, Abcam), as well as the corresponding control IgG antibodies [rabbit IgG (30000-0-AP, Proteintech) or mouse IgG (B900620, Proteintech)], were conjugated to the agarose beads. Protein lysate was then incubated with antibody-conjugated beads overnight at 4℃. After extensive washing with the kit’s wash buffer, bound proteins were eluted with elution buffer and prepared for Western blot analysis.

### Breast cancer mouse model

Breast cancer is a leading malignancy among young female patients who often experience POI and infertility following treatment with gonadotoxic anti-cancer agents (63, 64). We established a breast cancer mouse model to mimic the cancer-bearing conditions in young female cancer patients and investigate if pharmacological inhibition of JNK interferes with the anti-cancer efficacy of chemotherapy.

Eight-weeks-old C57BL/6 female mice were injected with mouse AT3OVA mammary carcinoma cell lines (SCC178, Millipore, USA), hereafter referred to as AT3 cells), following a previously established triple-negative breast cancer model protocol (65-67). In brief, AT3 cells were cultured in DMEM (2414543, Thermo Fisher Scientific) supplemented with 10% fetal bovine serum (FBS), 2 mM non-essential amino acids solution (11140050, Thermo Fisher Scientific), 15 mM HEPES (15630080, Thermo Fisher Scientific), 1xβ-mercaptoethanol (ES-007-E, Millipore, Burlington, MA, USA). Detached AT3 cells were centrifuged at 300 g for 3 min, and the cell pellets were resuspended with AT3 media mixed with Matrigel (354277, Corning, USA) at a concentration of 1X10^7^ cells/mL. To prevent Matrigel solidification, the cell suspension was loaded into a capped 1 mL syringe and kept on ice until injection. Mice under anesthesia were orthotopically injected with 50 μL of AT3 cell suspension into the fourth mammary fat pad. Tumor size was measured with a digital caliper and calculated using the formula: length width^2^ / 2= tumor size (mm^3^). Once the tumor reached ∼ 50 mm^3^, mice were randomly assigned to one of the following groups: vehicle control, chemotherapy (10 mg/kg DOXO or 150 mg/kg CPA through i.p.), co-administration of chemotherapeutic agent and JNK inhibitor SP600125, or JNK inhibitor alone.

For DOXO, breast cancer-bearing mice received 10 mg/kg DOXO via i.p. injection on days 0 and 7 (two doses in total). For CPA, mice were treated with a single dose of 150 mg/kg CPA via i.p. For co-administrated studies, 10 mg/kg JNK inhibitor SP600125 was administered via i.p. once, 2 h before the first dose of DOXO or CPA. All groups were monitored for survival and tumor growth throughout the treatment period. Ovaries were collected 14 days later for histology analysis and follicle counting.

### Statistics and reproducibility

All statistical analyses were performed using GraphPad Prism (version 5; GraphPad Software, San Diego, CA, USA). Data normality was assessed using the Shapiro-Wilk test before applying parametric tests. Unpaired Student’s t-tests were used to compare gene expression levels and the number of ovulated oocytes between two groups. One-way ANOVA was used to analyze differences in follicle counts, ovarian weight, vaginal opening, estrous cycle, and fertility outcomes between multiple groups. Changes in average litter size and tumor volume were evaluated using repeated measures two-way ANOVA, while two-way ANOVA assessed the effects of genotype and drug treatment on follicle counts. Tukey’s multiple comparison test was applied where correction for multiple comparisons was required. All fluorescence intensity, follicle counting and immunostaining experiments were independently performed with ≥ 3 biological replicates. Statistical significance was defined as *P* < 0.05.

## Data availability

The next generation sequencing datasets reported in this study are available from at the Gene Expression Omnibus (GSE314244).

## Authors’ roles

W. Zhao contributed to experimental design and execution, data collection, analysis and interpretation, and manuscript editing. J. Zhang and S. Liu contributed to bioinformatic analysis, data interpretation, and manuscript editing. Y. Wang contributed to initial conception, experimental design, and data collection and analysis. Y. Bo and MR. Choi contributed to data analysis, data interpretation, and manuscript writing and editing. Q. Zhang and S-Y. Kim contributed to manuscript editing. S. Xiao conceived of the project, designed experiments, analyzed and interpreted data, wrote the manuscript, and provided final approval of the manuscript.

## Funding and acknowledgments

This work was supported by the National Institutes of Health (NIH) R01HD115810 to S. Xiao and S-Y. Kim; K01ES030014 and P30ES005022 to S. Xiao; and Start-Up Fund from the Environmental and Occupational Health Sciences Institute (EOHSI) at Rutgers University to S. Xiao. This work was also partially supported by the Pilot Awards to S. Xiao from the Health and Environmental Science Institute (HESI, 2021514), Rutgers University Office of Research TechXpress Fund, and New Jersey Pediatric Hematology/Oncology Research Center of Excellence (NJ PHORCE).

