## Supplemental Figure S1 to S5 for "Inhibition of oogenic JNK preserves fertility and ovarian hormones during DNA-damaging cancer therapy"

### Slide 1
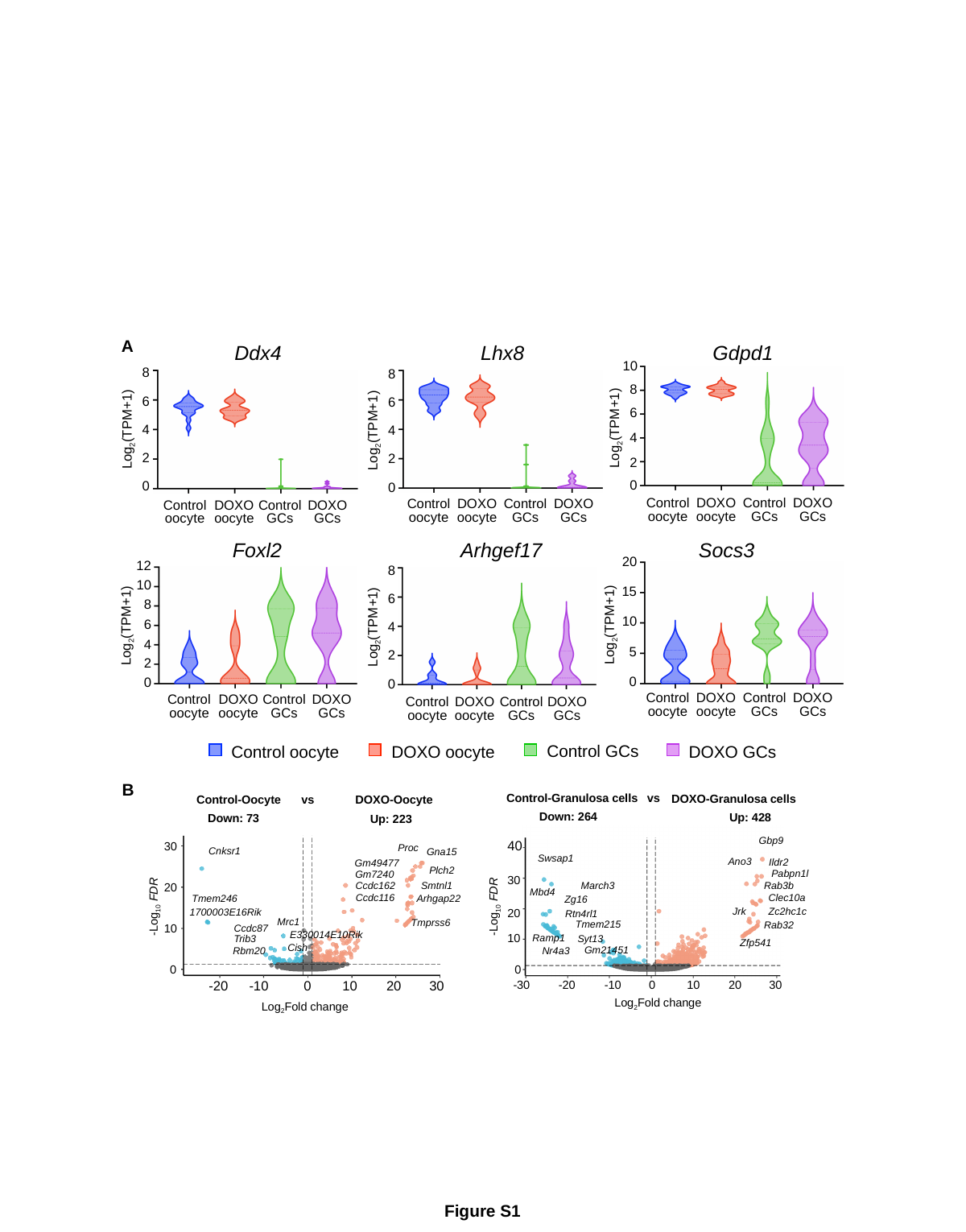

A
Ddx4
Lhx8
Gdpd1
10
8
6
Log_2_(TPM+1)
4
2
0
8
6
Log_2_(TPM+1)
4
2
0
8
6
Log_2_(TPM+1)
4
2
0
Control
oocyte
DOXO
oocyte
Control
GCs
DOXO
GCs
Control
oocyte
DOXO
oocyte
Control
GCs
DOXO
GCs
Control
oocyte
DOXO
oocyte
Control
GCs
DOXO
GCs
Foxl2
Arhgef17
Socs3
20
15
10
Log_2_(TPM+1)
5
0
12
8
6
Log_2_(TPM+1)
4
2
0
8
Log_2_(TPM+1)
6
4
2
0
10
Control
oocyte
DOXO
oocyte
Control
GCs
DOXO
GCs
Control
oocyte
DOXO
oocyte
Control
GCs
DOXO
GCs
Control
oocyte
DOXO
oocyte
Control
GCs
DOXO
GCs
Control GCs
DOXO oocyte
DOXO GCs
Control oocyte
B
Control-Granulosa cells
vs
DOXO-Granulosa cells
Control-Oocyte
vs
DOXO-Oocyte
Down: 264
Up: 428
Down: 73
Up: 223
Gbp9
40
30
Proc
Cnksr1
Gna15
Swsap1
Ano3
Ildr2
Gm49477
Plch2
Pabpn1l
Gm7240
30
Smtnl1
Rab3b
Ccdc162
March3
20
Mbd4
Clec10a
Ccdc116
Arhgap22
Tmem246
Zg16
-Log_10_ FDR
-Log_10_ FDR
Jrk
Zc2hc1c
1700003E16Rik
20
Rtn4rl1
Mrc1
Tmprss6
Tmem215
Rab32
10
Ccdc87
E330014E10Rik
10
Ramp1
Trib3
Syt13
Zfp541
Cish
Gm21451
Rbm20
Nr4a3
0
0
-20
-10
0
10
20
30
-30
-20
-10
0
10
20
30
Log_2_Fold change
Log_2_Fold change
Figure S1

### Slide 2
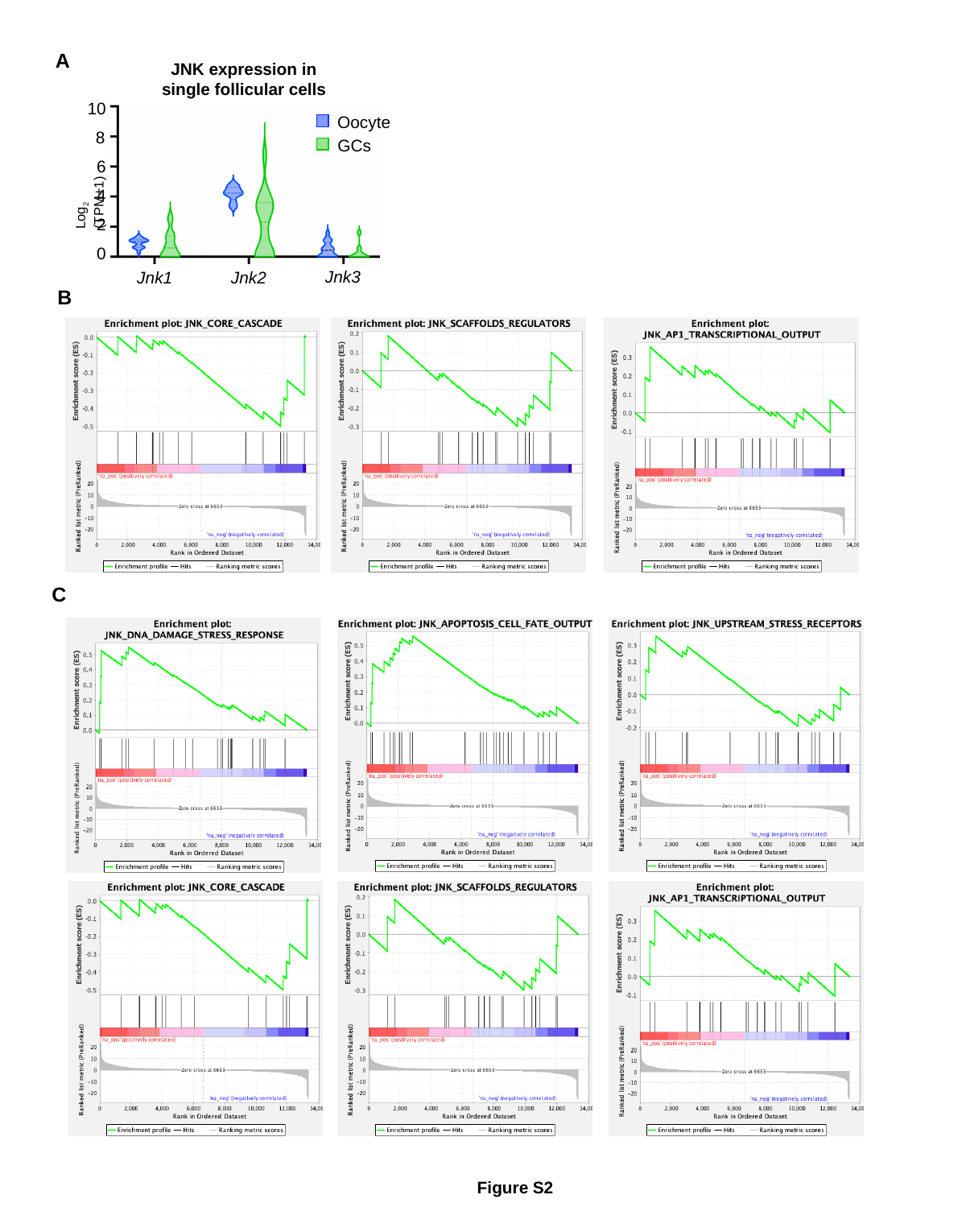

A
JNK expression in single follicular cells
10
Oocyte
8
GCs
6
Log_2_ (TPM+1)
4
2
0
Jnk3
Jnk1
Jnk2
B
C
Figure S2

### Slide 3
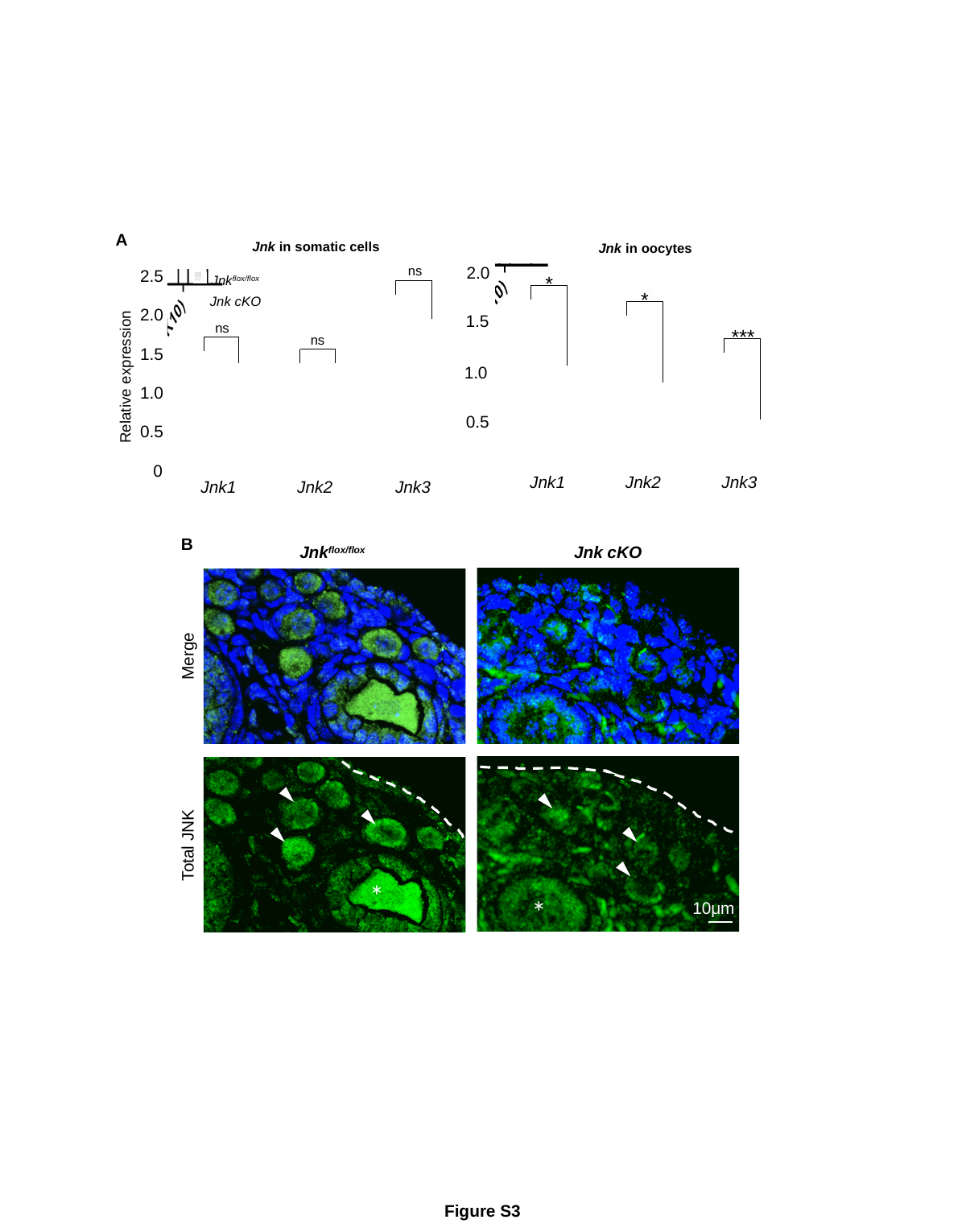

A
Jnk in somatic cells
Jnk in oocytes
2.0
ns
2.5
*
Jnk^flox/flox^
Jnk cKO
*
2.0
1.5
ns
***
ns
1.5
1.0
Relative expression
1.0
0.5
0.5
0
Jnk1
Jnk2
Jnk3
Jnk1
Jnk2
Jnk3
B
Jnk cKO
Jnk^flox/flox^
Merge
Total JNK
*
*
10μm
Figure S3

### Slide 4
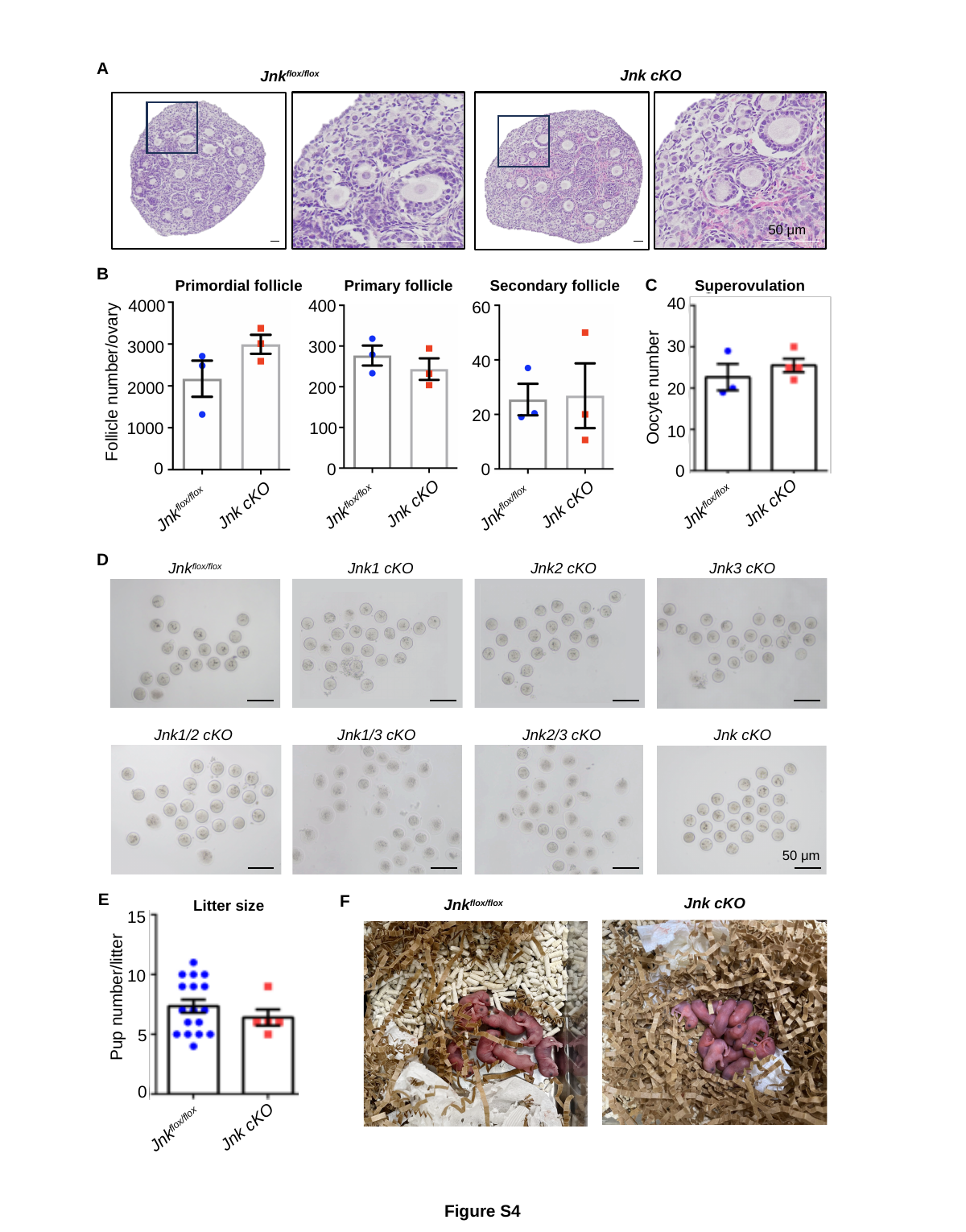

A
Jnk^flox/flox^
Jnk cKO
50 μm
B
C
Primordial follicle
Primary follicle
Secondary follicle
Superovulation
40
4000
400
60
30
3000
300
40
Follicle number/ovary
Oocyte number
2000
200
20
20
1000
100
10
 Jnk^flox/flox^
 Jnk cKO
 Jnk^flox/flox^
 Jnk cKO
 Jnk^flox/flox^
 Jnk cKO
0
 Jnk^flox/flox^
 Jnk cKO
0
0
0
D
Jnk^flox/flox^
Jnk1 cKO
Jnk2 cKO
Jnk3 cKO
Jnk1/2 cKO
Jnk1/3 cKO
Jnk2/3 cKO
Jnk cKO
50 μm
E
F
Jnk^flox/flox^
Jnk cKO
Litter size
15
10
Pup number/litter
5
0
 Jnk cKO
 Jnk^flox/flox^
Figure S4

### Slide 5
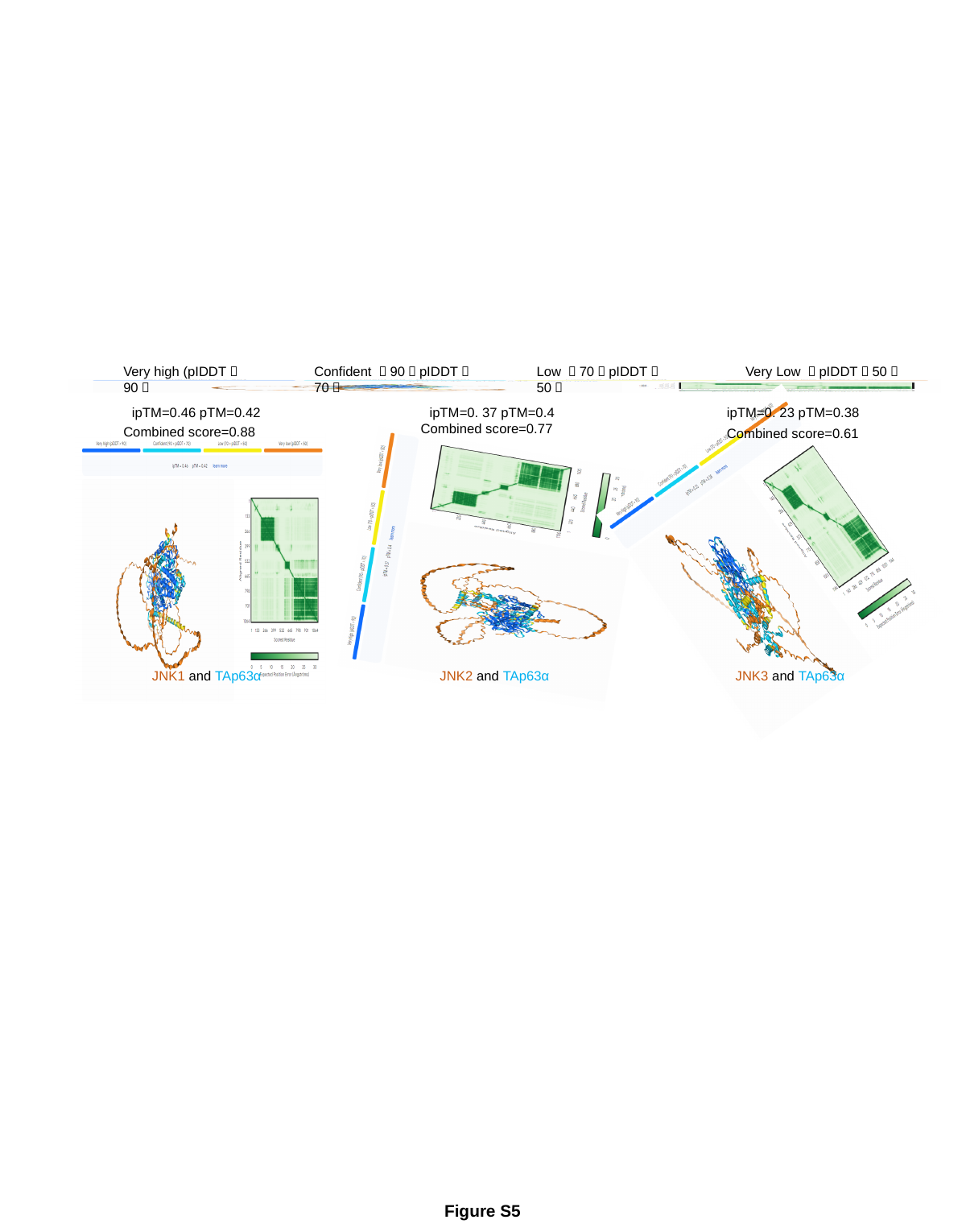

Very high (pIDDT＞90）
Confident （90＞pIDDT＞70）
Low （70＞pIDDT＞50）
Very Low （pIDDT＜50）
ipTM=0.46 pTM=0.42
ipTM=0. 37 pTM=0.4
ipTM=0. 23 pTM=0.38
Combined score=0.77
Combined score=0.88
Combined score=0.61
JNK1 and TAp63α
JNK2 and TAp63α
JNK3 and TAp63α
Figure S5
