## Supplemental Figure Legends for "Inhibition of oogenic JNK preserves fertility and ovarian hormones during DNA-damaging cancer therapy"

**Oocyte-intrinsic JNK signaling drives primary ovarian insufficiency and infertility induced by DNA-damaging anti-cancer agents**

Wenlong Zhao^1,2,3^, Jiyang Zhang^1,2^, Yingnan Bo^1,2^, Yingzheng Wang^1,2,4^, Mi Ran Choi^1,2^, Shichao Liu^1,2^, Qiang Zhang^5^, So-Youn Kim^6^, Shuo Xiao^1,2*^

^1^Department of Pharmacology and Toxicology, Ernest Mario School of Pharmacy, Rutgers University, Piscataway, NJ 08854, USA; ^2^Environmental and Occupational Health Sciences Institute (EOHSI), Rutgers University, Piscataway, NJ 08854, USA; ^3^Current address: Hangzhou Women's Hospital, Hangzhou Normal University, Hangzhou, Zhejiang 310008, China; ^4^Weill Institute for Cell and Molecular Biology, Cornell University, Ithaca, NY, 14853, USA; ^5^Gangarosa Department of Environmental Health, Rollins School of Public Health, Emory University, Atlanta GA 30322, USA; ^6^Department of Obstetrics, Gynecology and Reproductive Health, New Jersey Medical School (NJMS), Rutgers University, Newark, NJ 07103, USA

**Supplemental Figures and Figure legends**

**Figure S1.** **SMART-seq2 analysis reveals distinct transcriptomic profiles of oocytes and somatic cells following DOXO treatment.** **(A)** Violin plots showing normalized expression levels (log_2_[TPM+1]) of established marker genes for oocytes and granulosa cells in SMART-seq2 dataset. **(B)** Volcano plots of DEGs between control and DOXO-treated oocytes or granulosa cells, respectively (FDR＜0.05, fold change ＞2 or 0.5 compared to the controls).

**Figure S2. DOXO treatment induces transcriptomic changes featured by JNK-related signaling pathway in oocytes of primordial follicles.** **(A)** Expression of *Jnk1*, *Jnk2*, and *Jnk3* in oocytes and follicular somatic cells from single-cell SMART-seq2 dataset. **(B)** GSEA of oocyte gene profile on curated gene sets of JNK-related signaling pathways**. (C)** GSEA of granulosa cell gene profile on curated gene sets of JNK-related signaling pathways**.**

**Figure S3. Expression of JNK in *Jnk^flox/flox^* and *Jnk cKO* mouse ovaries**. **(A)** Relative mRNA expression of *Jnk1*, *Jnk2*, and *Jnk3* in follicular somatic cells (left) and oocytes (right) isolated from *Jnk^flox/flox^* and *Jnk cKO* mice, as determined by qRT–PCR (n=3 cells per group). Data are shown as mean ± SEM. Statistical significance was assessed using unpaired two-tailed Student’s t-test. ns, not significant, *p＞0.05*; *, *p < 0.05*; ***, *p < 0.001*. **(B)** Representative immunofluorescence images of total JNK (green) in ovarian sections from *Jnk^flox/flox^* and *Jnk cKO* mice. Oocyte of a primary follicle (asterisk), oocytes of primordial follicles (arrowheads) and ovarian cortex (dashed line) were indicated. Scale bar, 10 μm.

**Figure S4.** **Oocytes-specific deletion of JNK does not affect ovarian function and fertility.** **(A)** Representative histological images of ovarian sections from *Jnk^flox/flox^* and *Jnk cKO* mice at PND5. Scale bar, 50 μm. **(B)** Quantification of primordial, primary, and secondary follicles in *Jnk^flox/flox^* and *Jnk* cKO mice (n=3). **(C)** The count of ovulated oocytes in *Jnk^flox/flox^* and *Jnk* cKO mice under superovulation in *Jnk^flox/flox^* (n=3) and *Jnk cKO* (n=3) mice. **(D)** Representative images of superovulated oocytes from single and combined *Jnk* knockout models. Scale bar, 50 μm. **(E)** Litter size from *Jnk^flox/flox^* (n=17) and *Jnk cKO* mice (n=5). **(F)** Representative images of litters from *Jnk^flox/flox^* and *Jnk cKO* females. Data are presented as the mean ± SEM. Statistical significance was determined using an unpaired two-tailed Student’s t-test. ns, *p＞0.05*; *, *p＜0.05*; **, *p＜0.01*; ***, *p＜0.001*.

**Figure S5.** **AlphaFold3 prediction of the direct interaction between JNK1/2/3 and TAp63α.** Predicted 3D complex structures of JNK1, JNK2, or JNK3 with TAp63α generated by AlphaFold3. A combined ipTM + pTM score greater than 0.5 indicates a high likelihood of interaction. JNK1 and TAp63α interaction ipTM+ pTM=0.88; JNK2 and TAp63α interaction ipTM+ pTM=0.77; JNK3 and TAp63α interaction ipTM+ pTM=0.61.
